# Oral citrate supplementation mitigates age-associated pathological intervertebral disc calcification in LG/J mice

**DOI:** 10.1101/2024.07.17.604008

**Authors:** Olivia K. Ottone, Jorge J. Mundo, Boahen N. Kwakye, Amber Slaweski, John A. Collins, Qinglin Wu, Margery A. Connelly, Fatemeh Niaziorimi, Koen van de Wetering, Makarand V. Risbud

## Abstract

Despite the high prevalence of age-dependent intervertebral disc calcification, there is a glaring lack of treatment options for this debilitating pathology. Here, we investigate the efficacy of long-term oral K_3_Citrate supplementation in ameliorating disc calcification in LG/J mice, a model of spontaneous age-associated disc calcification. K_3_Citrate successfully reduced the incidence of disc calcification in LG/J mice without deleterious effects on vertebral bone structure, plasma chemistry, and locomotion. Notably, a positive effect on grip strength was evident in treated mice. Spectroscopic investigation of the persisting calcified nodules indicated K_3_Citrate did not alter the mineral composition and revealed that reactivation of an endochondral differentiation program in endplates may drive LG/J disc calcification. Importantly, K_3_Citrate reduced calcification incidence without altering the pathological endplate chondrocyte hypertrophy, suggesting mitigation of disc calcification primarily occurred through Ca^2+^ chelation, a conclusion supported by chondrogenic differentiation and Seahorse metabolic assays. Overall, this study underscores the therapeutic potential of K_3_Citrate as a systemic intervention strategy for disc calcification.

**Teaser:** Oral citrate mitigates intervertebral disc mineralization in a mouse model of age-dependent spontaneous disc calcification.

## Introduction

Intervertebral disc degeneration is a heterogeneous pathology linked to chronic low back and neck pain, which are consistently ranked among the leading causes of years lived with disability(1,2). Among the major phenotypes of disc degeneration, calcification is the least studied and understood(3,4). In humans, increased incidence of disc calcification is associated with aging, abnormal loading, and higher grades of degeneration and may occur in the nucleus pulposus (NP), annulus fibrosus (AF), or endplates (EP) of the disc, with or without other disc or spinal pathologies(5–8). With the increasing average human lifespan, age-associated disc calcification is of particular concern due to its association with pain and restricted range of motion(5,9,10).

Studies of human disc tissues and various animal models have shown that similarly to other soft tissues, calcification may be dystrophic or heterotopic in nature(11). Dystrophic calcification is characterized by amorphous calcium phosphate not associated with collagen but with a high phosphate-to-protein ratio and is thought to be caused by various cellular stressors that may disrupt the calcium-phosphate balance. By contrast, heterotopic ossification (HO) primarily driven by endochondral processes results in pathologic bone formation(3,12). Nucleating events for either form of ectopic calcification may include genetic susceptibility; tissue injury; local inflammation; cell death that leads to the release of Ca^2+^; membrane disruption leading to Ca^2+^ release or concentration in the mitochondria; extracellular vesicles; or the disruption of pyrophosphate (PPi) metabolism(3,4,12–14).

To date, no widely accepted therapy targeting disc calcification exists. In one clinical report, Boleto et al. demonstrated reductions of ochronosis-related low back pain and calcium deposition after the patient received the IL-1ra anakinra(15). An area unexplored in the treatment of disc calcification is non-pharmacological and non-biologic agents, which have shown efficacy in other pathologic calcification disorders(16). Notably, in the contexts of vascular and renal calcification, vitamins K and D as well as the chelating agents EDTA and citrate have been shown to reduce dystrophic calcification without affecting tissue architecture(17–20).

Citrate offers a particularly promising strategy, as a growing body of work demonstrates the importance of citrate in the maintenance of musculoskeletal tissues. For example, in humans, an oral K_3_Citrate supplement improves bone mineral density (BMD), without adverse effects(21,22). Subsequent studies established that citrate reduces bone loss through the inhibition of osteoclastogenesis(23,24). Moreover, a recent study showed that the citrate transporter SLC13a5 is central to the partitioning of citrate in mineralized tissues and essential to proper bone development(25). Notably, *ank*/*ank* mice with functionally deficient ANK, an ATP and citrate efflux channel, show extensive pathological mineralization of the spine and major articular joints, indicating a possible contribution of citrate, in addition to PPi, to the regulation of disc calcification(26,27). In addition to these beneficial effects on the skeleton, a recent preclinical study suggested that dietary citrate supplementation promotes ketogenesis, leading to improved longevity, metabolic health, and memory(28). These studies provide strong evidence of the safety of dietary citrate in multiple contexts and considering the known ability of citrate to chelate calcium, form the basis of our investigation.

We recently reported LG/J inbred mice as the first mouse model of spontaneous age-associated disc calcification(29). This disc calcification was associated with elevated free calcium and transcriptomic signatures relating to endochondral bone and calcium-phosphate homeostasis, with parallels to a subset of degenerated human NP tissues(29,30). Of note, LG/J mice are considered super-healers for their ability to heal injuries to ear and articular cartilage (31–34). Interestingly, in response to the destabilization of medial meniscus (DMM) injury, young LG/J mice develop robust ectopic calcification of the meniscus and synovium, suggesting calcification as a sequelae of the repair process (35). We, therefore, hypothesized that long-term oral K_3_Citrate supplementation would slow the age-dependent progression of disc calcification in LG/J mice through calcium chelation and by modifying the differentiation and/or metabolism of mineralizing cells. We discovered that K_3_Citrate supplementation effectively reduces disc calcification as well as attenuates age-associated meniscal and synovial calcification in LG/J mice. Our results suggest this mitigation occurs through calcium chelation, without impacting the underlying cellular processes driving calcification. Importantly this is the first study to demonstrate the ability of a widely available dietary supplement to disrupt age-associated disc calcification, offering a promising glimpse into citrate as a possible therapy.

## Results

### Long-term K_3_Citrate supplementation reduces age-associated disc calcification in LG/J mice without adverse systemic effects

Between 18 and 23 months of age, LG/J mice develop robust intervertebral disc calcification in the caudal spine, showing a strong dependence of phenotype on spine aging(29). To investigate the therapeutic potential of citrate to ameliorate disc calcification, we provided LG/J mice animals with 80 mM K_3_Citrate through drinking water from 17 months of age (prior to the development of calcification) until euthanasia at 23 months-of-age (Fig. 1a) (20,36). *In vivo* μCT analysis conducted at 22 months of age (Fig. 1B), revealed a significant reduction in the incidence of disc calcification in the K_3_Citrate-treated mice (Fig. 1C-D’). Notably, behavioral assays evidenced an increase in grip strength, an important metric used to assess frailty in humans (Fig. E, E’), without any changes in open field test, suggesting maintenance of ambulation in K_3_Citrate treated mice (Fig. F, F’)(37).

**Figure 1.**
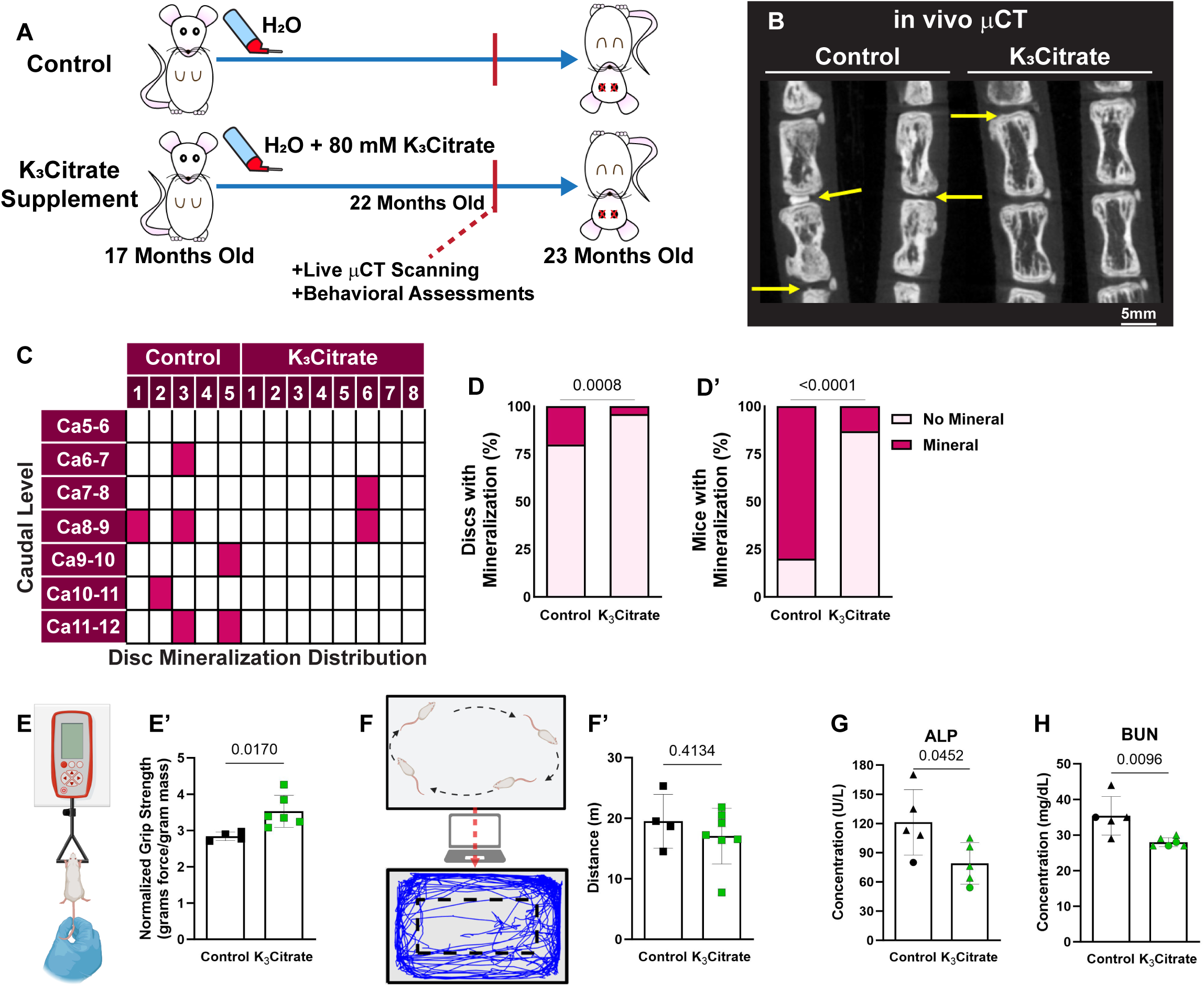
in vivo μCT shows K_3_Citrate supplementation improves ectopic calcification outcomes. **(A)** LG/J mice in the Control group and K_3_Citrate group received either regular drinking water or water continuously supplemented with 80mM of K_3_Citrate from 17 months-of-age until euthanasia at 23 months-of-age. **(B)** in vivo μCT demonstrated substantially **(C)** reduced incidence of disc mineralization, with respect to the **(D)** proportion of mineralized discs and **(D’)** proportion of mice with mineralized discs (Control: n=5 mice (2F, 3M); K_3_Citrate: n=8 mice (3F, 5M)). **(E-E’)** K_3_Citrate mice demonstrated higher grip strength than Controls (Control: 4 mice (2F, 2M); K_3_Citrate: n=6 mice (3F, 3M)). **(F-F’)** Open field analysis showed no differences in mobility in the K_3_Citrate cohort (Control: n=4 mice (2F, 2M); K_3_Citrate: n=7 mice (3F, 4M)). Slight reductions in plasma Alp (**G)** and BUN **(H)** were observed (Control: n=5 mice (1F, 4M); K_3_Citrate: n=5-6 mice (1-2F, 4)). Data are shown as mean ± SD. Distribution statistics were determined using a χ^2^ test. Behavioral and plasma statistics were determined using an unpaired t-test or Mann-Whitney test, as appropriate.

We then performed plasma analyses to determine systemic effects of K_3_Citrate supplementation. While the tissue non-specific alkaline phosphatase (ALP) (Fig. 1G) and blood urea nitrogen (BUN) (Fig. 1H) were lower in the K_3_Citrate-treated cohort, they were within physiological ranges reported in mice(29,38). Plasma albumin, calcium, chloride, glucose, phosphorus, and the calcification inhibitor fetuin-A remained unchanged by the treatment (Suppl. Fig. 1A-F). Similarly, the mouse inflammation marker GlycA did not change with K_3_Citrate (Suppl. Fig. 1G). Additionally, indicators of metabolic regulation: protein, total branched chain amino acids (BCAA), leucine, isoleucine, valine, alanine, acetoacetate, acetone, total ketone bodies, βhydroxybutyrate, chelatable magnesium (Mg^2+^), citrate, ApoA-1, ApoB, total triglyceride, total cholesterol, total calibrated low-density lipoprotein particle (cLDLP), and total calibrated high-density lipoprotein particle (cHDLP) also remained stable with K_3_Citrate supplementation (Suppl. Fig. 1H-Y). In conclusion, these extensive plasma analyses did not reveal any adverse effects of long-term K_3_Citrate supplementation.

Following euthanasia at 23 months, *ex vivo* μCT was conducted to further evaluate calcification nodules and vertebral structure. 2-dimensional planar views and 3-dimensional reconstructions of spinal motion segments showed disc calcification in control and K_3_Citrate cohorts (Fig. 2A, A’). While the incidence of disc calcification was higher than observed with *in vivo* μCT scanning one month prior, this is likely a reflection of the progressive pathology and the scanning resolution (see methods), and, showed marked reductions in the proportion of mineralized discs (Fig. 2B) size distributions of disc calcification (Fig. 2B’), calcification volume (Fig. 2C, C’), calcification density (Fig. 2D), and disc height (Fig. 2E) in K_3_Citrate treated mice, confirming the efficacy of K_3_Citrate supplementation in reducing disc calcification burden.

**Figure 2.**
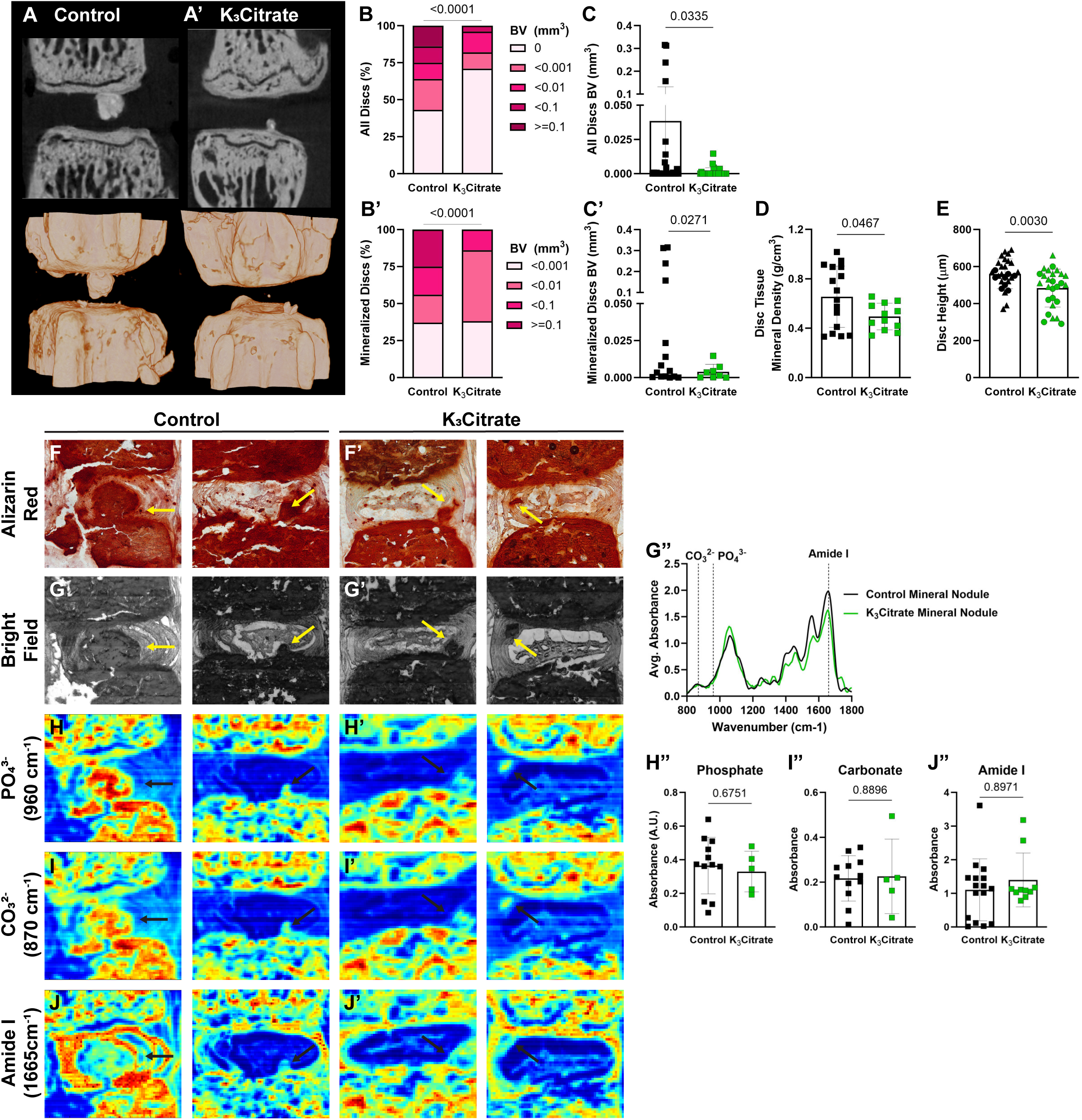
ex vivo μCT reveals quantitative alterations in mineral nodule incidence of K_3_Citrate mice, without changes to nodule composition. **(A-A’)** 2-D images and 3-D reconstructions show reductions in the **(B, C)** incidence and **(B’, C’)** size of disc mineralization in K_3_Citrate-treated LG/J mice. **(D)** K_3_Citrate supplementation resulted in lower mineral density in LG/J calcification nodules. **(E)** disc height decreased with K_3_Citrate supplementation. (Control mice: n=7 mice (2F, 5M); K_3_Citrate mice: n=7 mice (3F, 4M); 2 vertebrae/mouse; 28 discs, 14 vertebrae/treatment) Data are shown as mean ± SD. Significance was determined using an unpaired t-test or Mann-Whitney test, as appropriate. Distribution statistics were determined using a χ^2^ test. **(F-F’)** Alizarin red staining shows free calcium in LG/J discs is restricted to mineralized tissues **(G-G’)** FTIR bright field image scans showing mineral nodules in Control and K_3_Citrate mice, and **(G”)** normalized absorbance spectra reflect alignment of chemical composition across treatment conditions. **(H-J”)** Chemical maps of at **(H-H”)** phosphate (PO_43-_, 960 cm^-1^), **(I-I”)** carbonate (CO_32-_, 870 cm^-1^), and **(J-J”)** amide I (1665 cm^-1^) peaks reveal no changes in calcification nodule composition in K_3_Citrate mice. (Control mice: n=7 mice (2F, 5M); K_3_Citrate mice: n=7 mice (3F, 4M); 2 discs/mouse, 14 discs/treatment; Ca8-Ca10) Data are shown as mean ± SD. Significance was determined using an unpaired t-test or Mann-Whitney test, as appropriate.

To further examine the mineralized nodules in LG/J discs, Alizarin Red staining was conducted (Fig. 2F-F’), showing a concurrent abundance of calcium with the presence of mineralized nodules (29). Fourier transfer infrared (FTIR) spectroscopy was then used to evaluate if K_3_Citrate impacted the mineral composition. From these scans, brightfield images (Fig. 2G-G’) were used to identify mineral nodules in both treatment cohorts, and the averaged spectra were analyzed at absorbance peaks for phosphate (960 cm^-1^) (Fig. 5H-H”), carbonate (870 cm^-1^) (Fig. 5I-I”), and amide I (1665 cm^-1^) (Fig. 5J-J”). For all measured peaks, no differences were observed, as reflected in average absorbance curves for control and K_3_Citrate nodules (Fig.5G”).

### K_3_Citrate supplementation mitigates disc calcification without major structural or compositional impacts on NP and AF compartments

Safranin O/Fast Green/Hematoxylin staining was performed to evaluate disc morphology in treated mice (Fig. 3A, A’). Modified Thompson grading (Fig. 4 B, B’) did not show morphological changes with K_3_Citrate supplementation, suggesting that degeneration of the NP and AF compartments was not driven by disc calcification. In support of this, quantitative immunostaining for the NP phenotypic marker carbonic anhydrase 3 (CA3) (Fig. 3C-C”) showed no changes with K_3_Citrate supplementation. However, abundance of glucose transporter 1 (GLUT 1) (Fig. 3D-D”) was higher in the NP of K_3_Citrate-treated mice, suggesting that reduction in disc calcification preserves NP cell metabolism during aging. Picrosirius red staining was then used to assess collagen fiber thickness across cohorts. Bright field images (Fig. 3E, E’) demonstrated fibrotic remodeling of the NP in both cohorts (Fig. 3F), and quantitative polarized light imaging (Fig. 3G, H’) showed no differences in collagen fiber thickness in the NP, AF, or EP between vehicle and K_3_Citrate treated mice (Fig. 3H).

**Figure 3.**
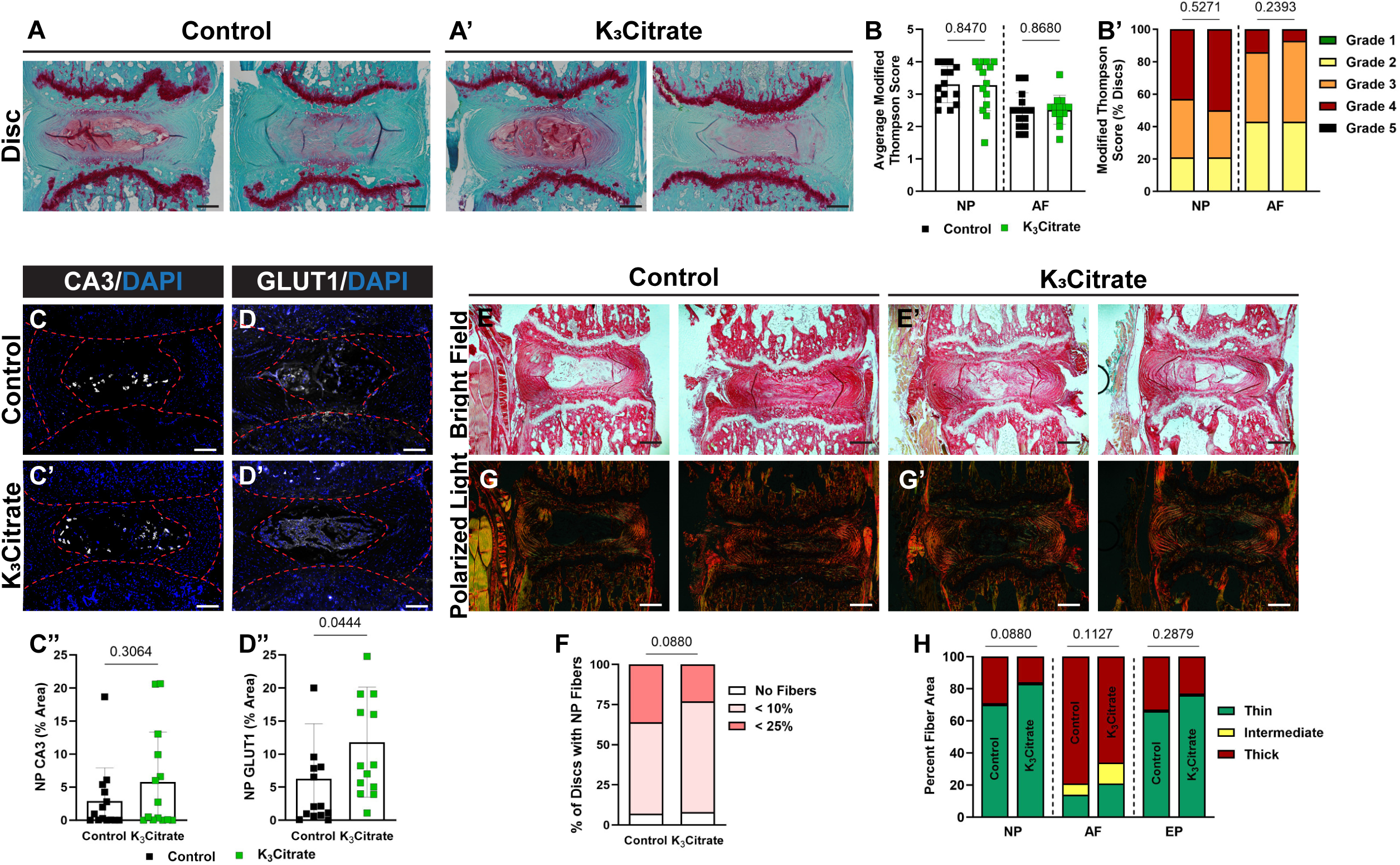
Quantitative histology reveals limited alterations to disc structure and cellular phenotype with K_3_Citrate supplementation. **(A-A’)** Representative Safranin O/Fast Green/Hematoxylin-stained discs, showing the range of mild and severe degeneration in LG/J Control and K_3_Citrate mice. Grading assessment using the **(B-B’)** modified Thompson scale to assess the NP and AF demonstrated no change to disc structure in K_3_Citrate mice. Abundance of NP phenotypic marker **(C-C’)** carbonic anhydrase (CA3) did not change with K_3_Citrate supplementation, but **(D-D’)** glucose transporter 1 (GLUT1) was more abundant in K_3_Citrate mice. **(E-E’)** Picrosirius red staining imaged in the bright field showed **(F)** no changes to the incidence of NP fibrosis with K_3_Citrate supplementation. **(G-G’)** Visualization under polarized light **(H)** showed no changes to collagen fiber thickness in the NP, AF, or EP of K_3_Citrate mice. (Control mice: n=7 mice (2F, 5M); K_3_Citrate mice: n=7 mice (3F, 4M); 2 discs/mouse, 14 discs/treatment; Ca6-Ca8) Data are shown as mean ± SD. Distribution statistics were determined using a χ^2^ test.

**Figure 4.**
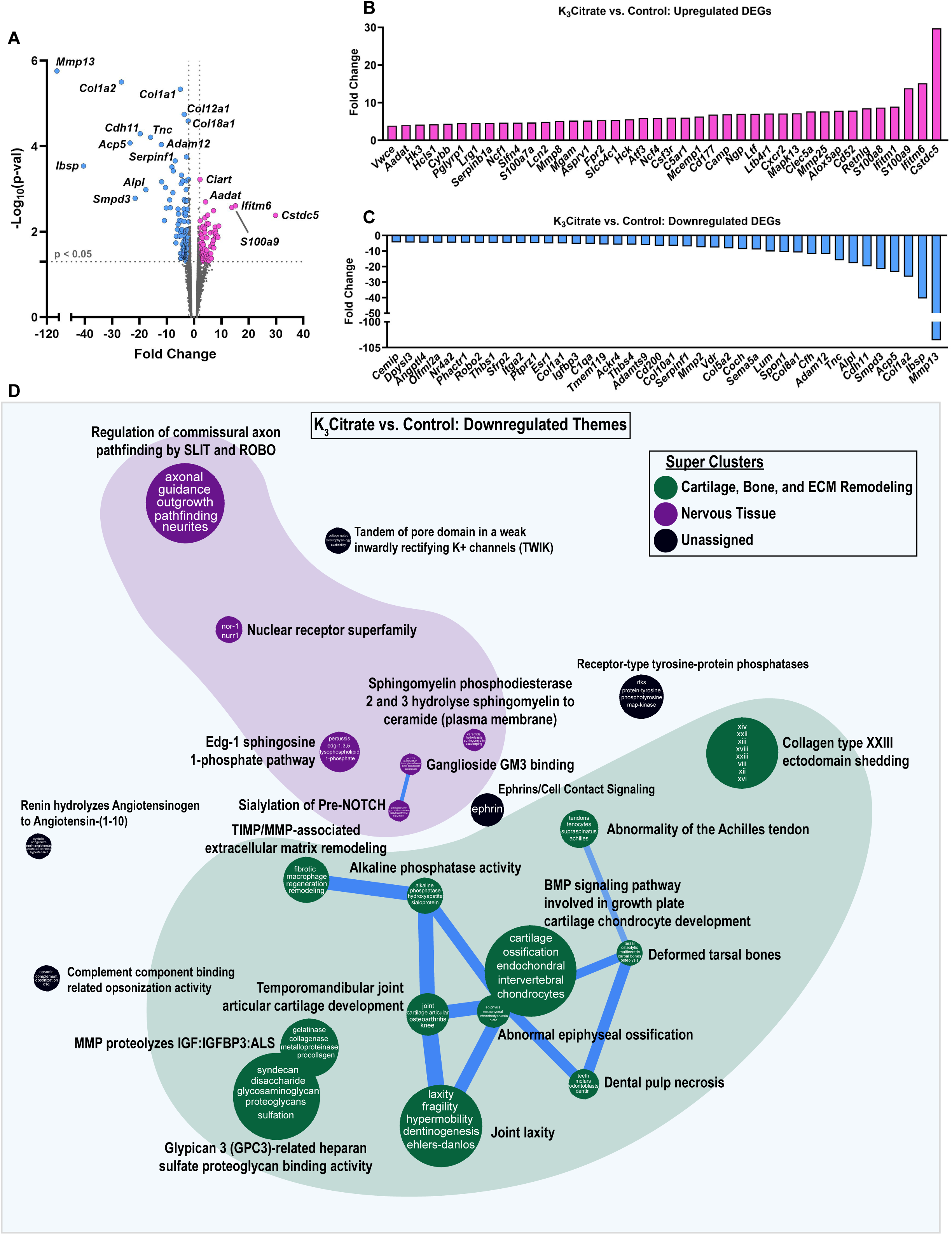
RNA-sequencing of NP tissues shows K_3_Citrate dampens signatures associated with cartilage and bone. **(A)** Volcano plot showing differentially expressed genes (DEGs) in NP tissues from control and K_3_Citrate-treated LG/J mice; *Mmp13*, *Col1a2*, and *Col1a1* are the most significant DEGs. **(B)** 39 DEGs were upregulated, and **(C)** 42 DEGs were downregulated. **(D)** Pathway-level thematic enrichment analysis conducted in CompBio highlighted thematic super clusters for *Cartilage, Bone, and ECM remodeling* (green) and *Nervous Tissue Development* (purple). (Control: n=4 mice (1F, 3M); K_3_Citrate: n=7 mice (2F, 2M))

To gain further insights into how K_3_Citrate supplementation and a reduction in disc calcification may have impacted the behavior of NP cells, we conducted RNA-seq analysis on NP tissues. Across treatment cohorts, 216 genes were differentially expressed (DEGs) (p<0.05, fold-change>2) (Fig. 4A), showing 86 upregulated (Fig. 4B) and 130 downregulated (Fig. 4C) DEGs. Pathway-level analysis was then conducted on upregulated and downregulated DEGs using the CompBio (PercayAI Inc., St. Louis, MO) tool to determine thematic associations among these genes. While upregulated DEGs demonstrated a weaker thematic enrichment than the downregulated DEGs (Suppl. Fig. 2A, B), many of these themes coalesced around a signal for *Immune/Inflammatory Process* or *Metabolism*. One of the metabolic themes was *Fructose-bisphosphate aldolase activity*, which could indicate increased glycolysis in the K_3_Citrate-treated cohort, aligning with higher GLUT1 abundance (Suppl. Fig. 2B).

Interestingly, analysis of the downregulated DEGs showed strong enrichment around *Cartilage, Bone, and ECM Remodeling*, and *Nervous Tissue* (Fig. 4D). The strongest gene signals within each thematic super cluster were: *Mafb*, *Col1a2*, *Sdc1*, *Col12a1, Col5a1*, *Col3a1*, and *Col10a1* (*Cartilage, Bone, and ECM remodeling*); and *Nr4a2*, *S1pr1*, *St8sia1*, *Smpd3*, and *Robo2* (*Nervous Tissue*) (Suppl. Fig. 2A’). This signature suggests that K_3_Citrate-mediated reduction in disc mineralization in LG/J mice likely limits the dedifferentiation of NP cells toward a chondrogenic phenotype. Though major structural differences beyond the reduction of calcification were not evident between cohorts, these findings do provide evidence of mild changes to the NP cell phenotype due to reduced mineralization.

### K_3_Citrate does not disrupt endochondral remodeling of the endplates

Although K_3_Citrate supplementation did not alter the morphology of NP or AF compartments in LG/J discs, SafO/Fast Green staining showed hypertrophic chondrocytes in what appeared to be a robust endochondral remodeling of the endplates in both control and K_3_Citrate-treated cohorts (Fig. 5A, A’)(29). Interestingly, the area of endochondral masses did not change with treatment, suggesting that K_3_Citrate did not alter the cellular processes driving calcification (Fig. 4B). Additionally, the chondrocytes showed robust aggrecan (ACAN) (Fig. 5 C-C”) and collagen X (COLX) (Fig. 5 D-D”) expression, providing molecular evidence that endplate cells were undergoing hypertrophic differentiation. TUNEL staining evidenced apoptosis in the bony endplates (Fig. 5 E-E’), however, there was no difference in cellularity (Fig. 5 E”) or fraction of TUNEL-positive cells (Fig. 5 E’”) between cohorts suggesting unhindered differentiation and maturation of chondrocytes. Considering that LG/J is a super healer strain and in conjunction with their propensity for mineralization in response to injury, these results suggest that intervertebral disc calcification in LG/J mice may be in part driven by robust endochondral healing response. This healing is likely a response to accumulated injury in the bony endplate with aging, wherein osteochondroprogenitor cells initiate a repair response that results in a calcified callus and subsequent propagation of the calcified nodules in the disc(32–35,39,40).

**Figure 5.**
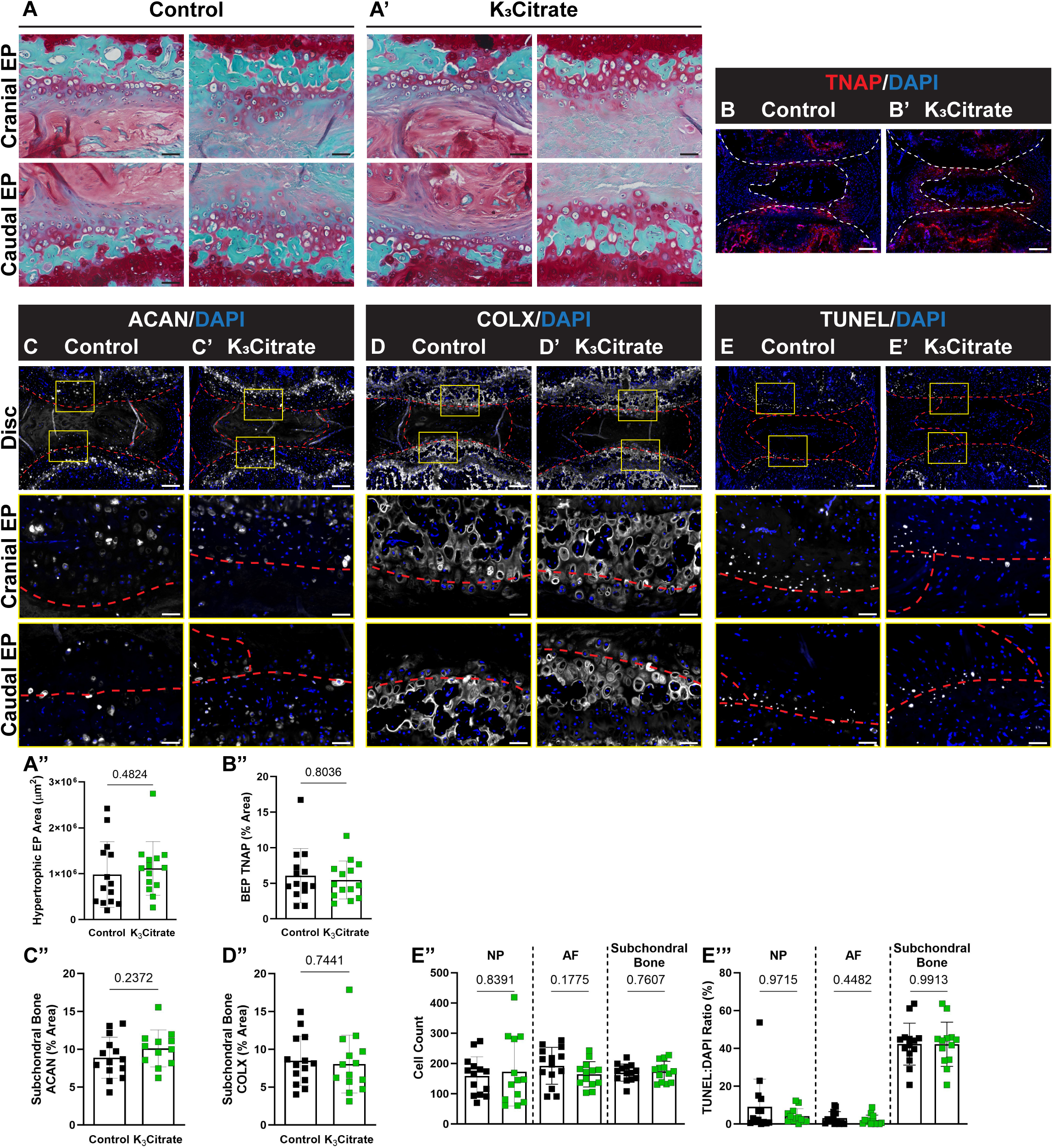
Quantitative histology reveals endplate chondrocytes and a chronic repair response may drive disc mineralization in LG/J mice, without a cellular response to K_3_Citrate. (**A-A”)** Safranin O/Fast Green/Hematoxylin-staining revealed what appeared to be aggregates of hypertrophic chondrocytes in the cartilaginous endplates, and the (**B)** area of these aggregates did not change with K_3_Citrate supplementation. Quantitative immunohistological staining subchondral bone/endplate space for hypertrophic chondrocyte markers (**C-C”)** aggrecan (ACAN) and (**D-D”)** collagen X (COLX), as well as **(E-E”’)** TUNEL staining to delineate cell death, provide evidence of lesions along the cartilaginous endplates, resembling fracture healing in bone; this was unattenuated in K_3_Citrate mice. (Control mice: n=7 mice (2F, 5M); K_3_Citrate mice: n=7 mice (3F, 4M); 2 discs/mouse, 14 discs/treatment; Ca6-Ca8) Data are shown as mean ± SD. Significance was determined using an unpaired t-test or Mann-Whitney test, as appropriate.

Together, these studies revealed three key findings: 1) disc calcification in LG/J mice appears in part to be driven by an endochondral remodeling process, driven by chondrocytes in the bony endplates; 2) fibrotic degeneration of the disc occurs independent of calcification status; and 3) K_3_Citrate supplementation effectively reduces the incidence of disc calcification, leading to alterations to the underlying cellular processes in the NP but not in the endplate.

### K_3_Citrate supplementation minimally impacts vertebral bone and knee joint structure in LG/J mice

Previous studies have shown that K_3_Citrate improves bone health in humans and mice; we therefore assessed the effect of treatment on vertebral bone morphology (21,22,24). Accordingly, 3D reconstructions of caudal vertebrae (Suppl. Fig. 3A, A’) were evaluated and showed no changes to vertebral length in K_3_Citrate mice (Suppl. Fig. 3B). Similarly, trabecular bone properties of BV/TV, trabecular separation (Tb. Sp.), Tb. Th., trabecular number (Tb. N.), and bone mineral density (Suppl. Fig. 3C-G) did not change with K_3_Citrate supplementation. However, evaluation of the cortical bone (Suppl. Fig. 3H, H’) showed mild cortical thinning, evidenced by lower bone volume (BV), tissue mineral density, cross-sectional thickness (Cs. Th.), and bone area (B. Ar.) without changes to the closed porosity or bone perimeter (B. Pm.) (Suppl. Fig. 3I-N) in K_3_Citrate mice. Treatment did not affect the plasma levels of IFN-γ, IL-1β, IL-2, IL-4, IL-5, IL-6, IL-10, IL-12/p70, IL-15, Il-17A/F, IL-27/p28/IL-30, IL-33, IP-10, KC/GRO, MCP-1, MIP-1a, MIP-2, and TNF-α (Suppl. Fig. 3O -FF), indicating cortical thinning was not the result of systemic inflammation. Importantly, the limited cortical thinning of approximately 5%, is unlikely to translate into altered bone function (41–43). Taken together, these results demonstrate the ability of oral K_3_Citrate supplementation to mitigate disc calcification without adverse systemic effects.

Prior to this investigation, no studies had investigated LG/J knees in the context of aging, with the only report on LG/J knee phenotypes being in 8-week-old animals in response to DMM injury (35). Interestingly, μCT analysis revealed significant synovial, meniscal, and patellar calcification in both control and K_3_Citrate cohorts (Suppl. Fig. 4A, A’). Quantification of the number of calcified nodules in the synovium (Suppl. Fig. 4B), meniscus (Suppl. Fig. 4C), and patella (Suppl. Fig. 4D) revealed that in the control mice, synovial and meniscal nodules were fewer in number but larger in size than in the K_3_Citrate cohort. This suggests that K_3_Citrate limited the development of calcification in the knees of LG/J mice. H&E (Suppl. Fig. 4E, E’) and Toluidine Blue (Suppl. Fig. 4F, F’) staining did not reveal differences in the overall structural integrity of the knee joints (Suppl. Fig. 4G-J) (44,45). Similarly, histomorphometric analysis of the articular cartilage (Suppl. Fig. 4K, K’), calcified cartilage (Suppl. Fig. 4L, L’), and subchondral bone (Suppl. Fig. 4M, M’) showed no changes between control and K_3_Citrate-treated knees, suggesting that despite robust calcification of the knee joint, articular cartilage in LG/J mice is not susceptible to age-associated osteoarthritis.

### K_3_Citrate supplementation reduces mineralization without altering the chondrogenic differentiation program and metabolism

To further investigate the hypothesis that K_3_Citrate limits endplate-mediated intervertebral disc calcification through the chelation of calcium, without impacting cellular processes, we used an *in vitro* model of endochondral differentiation. Since technical challenges prevent the culture of primary mouse endplate cells, the ATDC5 mouse cell line, which models endochondral ossification, transitioning from chondrogenic to osteoblastic differentiation under appropriate culture conditions was chosen(46–48). Accordingly, ATDC5 cell differentiation was studied in the presence of either 0.25 mM K_3_Citrate or 0.50 mM K_3_Citrate and mineralization, differentiation status, and metabolic processes were assessed in the proliferating (7-day), hypertrophic (14-day), and transition stage between hypertrophic chondrocytes and osteoblasts (21-days) (Fig.6A) (48).

**Figure 6.**
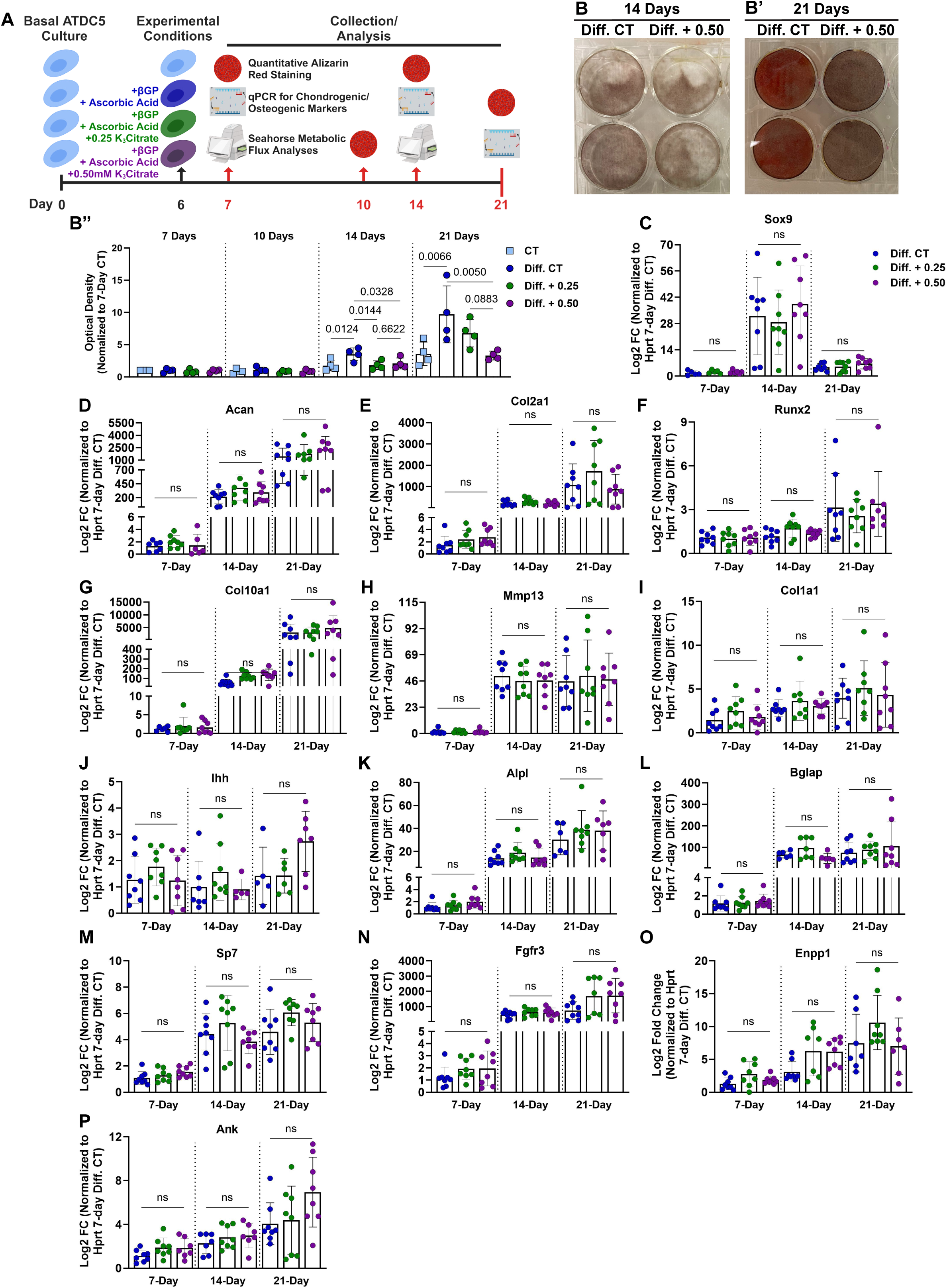
K_3_Citrate supplementation reduces mineralization without impacting the cell differentiation program in ATDC5 cells, used to model chondrogenic differentiation. **(A)** Schematic showing the experimental timeline and strategies used to understand how K_3_Citrate disrupts mineralization during chondrogenic differentiation. **(B-B”)** Representative images of Alizarin Red staining in differentiated control (Diff. CT) and differentiated ATDC5 cells treated with 0.50 mM K_3_Citrate (Diff. + 0.50) and quantification of all treatment groups shows a reduction in mineralization of ATDC5 cell cultures treated with K_3_Citrate. (n=4 sets/timepoint, 2 averaged replicates/set) mRNA evaluation of various markers of chondrogenic differentiation demonstrate that throughout differentiation, K_3_Citrate supplementation does not alter progression through this program: **(C)** Sox9, **(D)** Acan, **(E)** Col2a1, **(F)** Runx2, **(G)** Col10a1, **(H)** Mmp13, **(I)** Col1a1, **(J)** Ihh, **(K)** Alpl, **(L)** Bglap, **(M)** Sp7, **(N)** Fgfr3, **(O)** Ank, and **(P)** Enpp1. (n=8 sets/timepoint, 2 averaged replicates/set) Data are shown as mean ± SD. Significance was determined using an ANOVA or Kruskal-Wallis test, as appropriate.

Quantitative alizarin red staining showed increased mineralization in the differentiated control (Diff. CT) relative to the undifferentiated control (CT) by 14 days, and this was more pronounced by 21 days (Fig.6B-B”). Further, this increase in mineralization was reduced by treatment with 0.25 mM (Diff. + 0.25) and 0.50 mM (Diff. + 0.50) K_3_Citrate at both time points. We then evaluated the expression of markers for different stages of chondrogenic differentiation. First, the success of the differentiation experiment was confirmed by comparing the differentiated and undifferentiated control groups for Sox9- and Runx2-regulated genes and pyrophosphate regulators (Suppl. Fig. 5A-N). Temporal variation in the Diff. CT group indicated cells differentiated toward a hypertrophic stage by 14 days and that at 21 days, cells remained in a transition stage between hypertrophic chondrocytes and endochondral ossification. The impact of K_3_Citrate was then evaluated, showing no differences in the expression of *Sox9*, *Acan*, *Col2a1*, *Runx2*, *Col10a1*, *Mmp13*, *Col1a1*, *Ihh*, *Alpl*, *Bglap*, *Sp7*, and *Fgfr3* (Fig.6C-P) across treatment groups at both timepoints. This lack of change in gene expression profiles and the reductions in alizarin red staining with K_3_Citrate suggested that calcium chelation is the predominant mechanism of reduced calcification in the LG/J endplate callus.

Previous reports have indicated that oral citrate supplements can alter cell metabolism through the inhibition of glycolysis(28,49). Accordingly, Seahorse metabolic flux assays were conducted at the 7-day and 14-day time points to assess whether metabolic switching contributed to the reduction in ATDC5 mineralization. The impact of K_3_Citrate on glycolytic capacity was evaluated using methods described by Moorkerjee et al(50). OCR (Fig.7A, Suppl. Fig. 6A) and ECAR (Fig.7B, Suppl. Fig. 6B) were recorded under conditions described in the methods. These measurements were used to calculate the proton production rate (PPR) (Fig.7C, Suppl. Fig. 6C), which showed that K_3_Citrate did not impact the glycolytic capacity of ATDC5 cells. We then calculated glycolytic and oxidative ATP production rates following K_3_Citrate supplementation (51,52). Again, OCR (Fig.7D, Suppl. Fig. 6D) and ECAR (Fig.7E, Suppl. Fig. 6E) traces were not different across treatment groups. Accordingly, the computed glycolytic and oxidative ATP production rates showed no change with K_3_Citrate supplementation (Fig.7F, Suppl. Fig. 6F). The results of these assays were further validated using the well-documented Mito Stress test (53,54). Again, our results showed that cells cultured with K_3_Citrate did not experience changes to maximum, ATP-linked, or spare oxygen consumption capacity (Fig.7G-I, Suppl. Fig. 6G-I). Taken together, these findings indicate that extracellular K_3_Citrate does not alter the metabolic function of differentiating chondrocytes, and that K_3_Citrate reduces calcification in LG/J mice by Ca^2+^ chelation.

**Figure 7.**
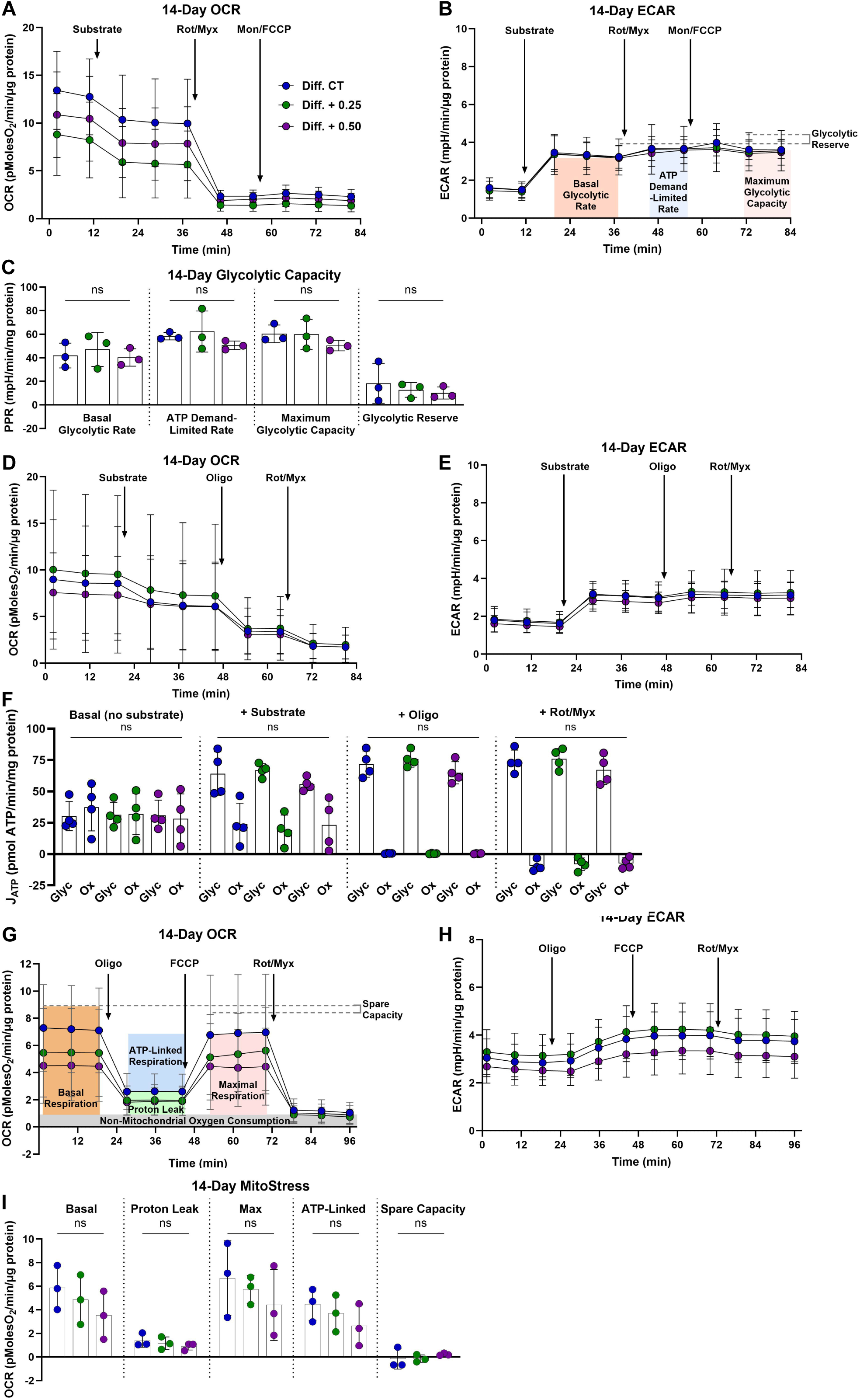
K_3_Citrate supplementation does not impact glycolytic or oxidative metabolism in ATDC5 cells cultured for 14 days. **(A)** OCR and **(B)** ECAR traces for ATDC5 cells cultured with or without K_3_Citrate **(C)** to evaluate glycolytic capacity and glycolytic reserve. **(D)** OCR and **(E)** ECAR traces for ATDC5 cells cultured with or without K_3_Citrate **(F)** to evaluate glycolytic and oxidative ATP production rates. **(G)** OCR and **(H)** ECAR traces for ATDC5 cells cultured with or without K_3_Citrate **(I)** for the classical Mito Stress test to evaluate key parameters of mitochondrial function. (n = 3 sets, 3-4 replicates/set) Data are shown as mean ± SD. Significance was determined using an ANOVA or Kruskal-Wallis test, as appropriate.

## Discussion

Intervertebral disc calcification is a prevalent subphenotype of age-dependent disc degeneration for which there is no current standard of care(5,55,56). Despite the negative impact of this phenotype on back pain and morbidity, the etiology of disc calcification is not well-established. Notably, studies delineating heterotopic and dystrophic calcification in the disc indicate multiple cellular mechanisms may govern the calcification process in a context-dependent manner(7,57). Historically, the study of disc calcification and the development of intervention strategies has been limited by a lack of mouse models which recapitulate this pathology, without the manipulation of a specific gene. Our group has previously described that LG/J, an inbred mouse strain, develops spontaneous age-associated caudal disc calcification, opening the door to new avenues of research(29). In this study, we show that a long-term oral K_3_Citrate supplementation successfully reduces the incidence of severity of age-associated, spontaneous disc calcifications in LG/J mice. Analyses of disc tissues in control and K_3_Citrate mice also highlighted that the calcification phenotype in LG/J mice is likely to be driven in part by endochondral processes originating in the endplates. Importantly, our studies suggest that K_3_Citrate supplementation reduces calcification through the chelation of excess calcium and does not interfere with the endochondral differentiation or cellular bioenergetics, cellular processes driving disc calcification.

Citrate was first identified as a physiologically relevant chelator of calcium in 1940 and has since been used in contexts of renal and vascular calcification to prevent the pathologic calcification(20,21,58). In musculoskeletal tissues, K_3_Citrate supplementation is shown to reduce osteoporotic outcomes by inhibiting osteoclastogenesis(21–23,59). Most notably, Pak et al. demonstrated K_3_Citrate supplementation reduced spinal bone loss in tandem with reducing kidney stones in patients being treated for calcium urolithiasis, demonstrating dual beneficial effects where citrate reduced dystrophic calcification while simultaneously preventing bone loss(21). While K_3_Citrate supplementation in mice has been shown to rescue osteopenic spinal phenotypes, the ability of K_3_Citrate to alter disc calcification had yet to be determined(24). Therefore, we tested the ability of K_3_Citrate to disrupt disc calcification in LG/J mice, a recently described model of spontaneous age-associated disc calcification(29). Remarkably, both *in vivo* and *ex vivo* μCT scans showed a significant reduction in disc calcification, highlighting the utility of K_3_Citrate in treating disc calcification.

At the systemic level, our results consistently demonstrated the safety profile of long-term K_3_Citrate supplementation. While behavioral analysis showed maintenance of the overall mobility of the K_3_Citrate mice, grip strength studies showed a small but consistent increase in the K_3_Citrate group. One possible explanation for this improvement is the supplementation of potassium, as lower potassium has been correlated to lower handgrip strength in older humans(60). It is also possible that the observed increase in grip strength results from increased intracellular citrate in muscle cells, though a specific study of the muscle in this model would be required to substantiate this hypothesis(61,62). Nevertheless, this finding highlights reduced frailty in treated mice. When plasma composition was analyzed, results indicated that plasma chemistry was not significantly altered by K_3_Citrate citrate, which is consistent with a previous report in humans(22). The two analytes that did change were TNAP and BUN, though both fell within previously reported physiological ranges for aging mice(29,38). Although ALP is broadly associated with PPi conversion to Pi and subsequent ectopic calcification, it was previously shown that systemic ALP levels are poor indicators of mineralization in LG/J mice(29). Regarding BUN, the reduction in the K_3_Citrate cohort could be indicative of a lower acid burden associated with treatment (36,38). In both cases, it is most likely that the observed decrease is not overtly significant in terms of its physiological consequence. Additionally, when caudal vertebrae were analyzed, there was no impact of K_3_Citrate supplementation on the structural properties of the trabecular bone; however, mild endocortical thinning was observed. Though this finding should not go unnoticed, considering the lack of change to trabecular bone and resilience of bone to small changes in bone volume, it is unlikely this cortical thinning manifested in reduced mechanical properties (41–43). Moreover, analyses of the knees demonstrated K_3_Citrate altered the joint calcification, without impact on the articular cartilage. Notably, these analyses also revealed that despite the robust calcification of LG/J knees, their articular cartilage is not susceptible to age-associated osteoarthritis, which could provide an interesting model for future comprehensive studies investigating the knee phenotype. Taken together, these findings generally support the safety of K_3_Citrate, showing minimal systemic effects while inhibiting ectopic joint calcification.

The original study identifying disc calcification in LG/J mice speculated the observed calcification was dystrophic in nature and may be the result of a combination of genetic predisposition, age-related stress, and tissue damage from cell death(29). This was supported by the enrichment of LG/J transcriptomic signatures related to calcium-phosphate homeostasis and cell death as well as a high phosphate to protein ratio in the mineral nodules (29). Analysis in the present study expands on these findings, showing that there may be an underlying endochondral and remodeling process involved in LG/J disc calcification. In support of this, RNA-sequencing analysis of NP tissues showed *Mmp13*, *Col1a2*, and *Col1a1* to be downregulated and the most significantly differentially expressed genes in the NP of K_3_Citrate mice. These are not only critical markers of fibrotic remodeling in the disc but also chondrogenic differentiation, providing evidence of the K_3_Citrate-driven reduction of disc calcification in LG/J mice could in part due to delayed NP cell differentiation toward a hypertrophic chondrocyte-like phenotype(63–66). Interestingly, abundant aggrecan, collagen 10, and robust safranin-o staining of the subchondral boney endplates in both LG/J cohorts provided evidence of a unique process involving re-activation of an endochondral differentiation contributing to disc calcification. This aligns closely with a previous study showing a transcriptomic signature related to endochondral bone and injury studies which demonstrated increased healing capacity in cartilaginous tissues of LG/J mice and a susceptibility to ectopic calcification in the presence of injury(29,31–35). In studies of ear puncture and full-thickness articular cartilage injury, LG/J mice are shown to fully resolve these injuries; and genetic studies correlated *Axin2, Wnt16, Xrcc2, and Pcna* with healing of both tissues, providing evidence that an enhanced DNA repair response and Wnt signaling are critical components of this unique wound healing(34). Interestingly, in response to DMM injury, LG/J mice develop robust synovial and meniscal calcification, correlated with SNPs relating to angiogenesis, bone metabolism/calcification, arthritis, and ankylosing-spondylitis and gene transcripts of *Aff3*, *Fam81a*, *Syn3*, and *Ank*(35). Correlating these observations with the disc calcification phenotype in LG/J mice, our findings suggest that disc calcification in LG/J mice may in part be due to an injury repair response to age-related wear of the bony endplates (39,40,67). It is known that endplate injuries are common, especially with aging, and may contribute to the degeneration of the NP and AF compartments(68,69). Accordingly, calcified cartilage along the CEP could serve as a nucleation site in the presence of cell death, which would align well with the mineralized nodules in LG/J discs ultimately being acellular, dystrophic calcifications and not as structured hydroxyapatite seen in bone(70). Importantly, when the composition of the mineralized nodules was analyzed, K_3_Citrate-treated mice did not differ from controls, indicating that while the size and quantity of the mineral nodules were greatly reduced, the end product formed was not chemically different because of K_3_Citrate. Together, these results suggested that K_3_Citrate was likely improving disc calcification outcomes through the chelation of calcium, without broadly impacting the underlying endochondral processes in the endplate.

To substantiate this hypothesis, we modeled mineralizing chondrocytes undergoing differentiation with the ATDC5 cells and found K_3_Citrate causing significantly reduced mineral deposition. Supporting the findings *in vivo*, the expression of genes controlling the progression of chondrogenic differentiation and calcification in ATDC5 cells was unchanged. There were also no changes in glycolytic or oxidative metabolism with K_3_Citrate supplementation; but our results did demonstrate an expected temporal switch toward oxidative metabolism between 7- and 14-day timepoints, which has previously been identified as an important feature of chondrogenic differentiation program in growth plate(53). This study clearly shows that K_3_Citrate supplementation safely and specifically targets ectopic calcification without modulating the underlying cellular and genetic causes.

It is well understood that the pathogenesis of disc calcification is multifactorial, which has complicated the development of intervention strategies. Among these factors, a proper balance of PPi metabolism has been linked to dystrophic calcification in the endplate and AF compartments of the disc, as shown in ANK and ENPP1 mutant mice (27,71,72). Of note, in the ANK model, transcriptomic analysis of disc tissues highlighted dysregulation of BMAL/CLOCK, underscoring the interplay of multiple complex processes regulating disc calcification. Studies of *Bmal1* show the importance of circadian regulation in disc health, with multiple knockout models leading to heterotopic calcification of the disc(73,74). Additionally, advanced glycation end products (AGEs), which are known to accumulate with aging, are associated with endochondral ossification of the disc, which provides insight into a possible mechanism to target in mediating disc calcification; but to date, this has not led to clinical interventions(75,76). Observations in scoliosis patients have also demonstrated the contribution of abnormal loading to CEP calcification, which can lead to more robust ectopic calcification impacting the NP, AF, or vertebrae (7,8,77). What these studies indisputably demonstrate is the complexity of disc calcification and the involvement of multiple processes in the onset of this pathology.

Excitingly, our work demonstrates the ability of K_3_Citrate – a low-cost dietary supplement – to intervene in the progression of disc calcification. Of significance, our results suggest the effect of K_3_Citrate is in large part through its known chemical properties as a calcium chelator, and, therefore, its beneficial effect is independent of the intricate cellular mechanisms driving disc calcification. While this leaves open many interesting scientific questions about the underlying biology of disc calcification, it also suggests that K_3_Citrate supplements could prevent or reduce disc calcification in a variety of disease contexts, due to its non-specific efficacy. Future studies should validate the ability of K_3_Citrate supplementation to mediate disc calcification in other animal models to more sufficiently confirm these findings and expand its applicability to human disease.

## Materials and Methods

### Mice, treatment, and study design

Animal procedures were performed under approved protocols by the IACUC of Thomas Jefferson University (TJU). LG/J mice (Stock #000675, Jackson Labs) were bred at TJU and aged to 23 months, when intervertebral disc mineralization occurs(29). Treatments for this study began when mice were 17 months old, prior to developing disc calcifications. All mice belonged to one of two treatment cohorts: control or K_3_Citrate. Mice in the control cohort received regular, untreated drinking water throughout the study. Mice in the K_3_Citrate cohort began receiving a continuous supplementation of 80 mM K_3_Citrate (Sigma-Aldrich, C3029) in their drinking water at 17 months-of-age. They received this continuous supplementation until the experiment’s conclusion. Mice were euthanized with CO_2_ asphyxiation.

All mouse experiments included male and female LG/J mice. Previous reports on the disc phenotype in LG/J mice show there are no sex-based differences, and this is a common finding in the mouse intervertebral disc(29,63,64,78).

### In Vivo Micro-Computed Tomography (μCT)

At 22 months-of-age, in vivo μCT scanning was conducted on the caudal regions of control (n=5) and K_3_Citrate (n=8) mice at the Small Animal Molecular Imaging Facility at TJU. Mice were anesthetized with 3% isoflurane. Once anesthetized, μCT scanning was conducted with an effective pixel size of 39.15 microns, field size of 40 mm by 35 mm, and exposure time of 30 minutes. Scans were visualized using Weasis DICOM Viewer (v4.0.3).

### Behavioral Tests

For all behavior tests, mice acclimated to the behavior testing room for one hour prior to testing. Forelimb grip strength of Control (n=4) and K_3_Citrate (n=6) mice was assessed using a Grip Strength Meter (DFIS-2 Series Digital Force Gauge, Columbus Instruments). To measure grip strength, animals held by their tails were allowed to tightly grasp a force gauge bar using both forepaws. Mice were then pulled away from the gauge until both limbs released the bar. Data recorded represents the average of five trials per mouse. Between trials, mice rested for one minute. An open field test was used to assess the general locomotion of Control (n=4) and K_3_Citrate (n=7) mice. In this test, mice were placed in an open field apparatus and recorded with an overhead camera for ten minutes. Video data were then processed in Matlab using the open-source code developed by Zhang et al(79) to determine the distance traveled by each mouse.

### Plasma Analyses

Blood was collected immediately postmortem by intracardiac puncture using heparinized needles. Plasma was separated from red blood cells via centrifugation at 1500 rcf and 4°C for 15 minutes and stored at -80°C until the time of analysis. Albumin, ALP, BUN, Calcium, Chloride, Glucose, and Phosphorus were analyzed using a custom blood chemistry panel (IDEXX BioAnalytics) (Control n=5, K_3_Citrate n =6). Fetuin-A was quantified using the mouse Fetuin-A/AHSG DuoSet ELISA (R&D Systems) according to the manufacturer’s instructions (Control n=7, K_3_Citrate n =7). Cytokine and proinflammatory marker concentrations were evaluated using the V-PLEX Mouse Cytokine 19-Plex Kit (Meso Scale Diagnostics, K15255D) according to the manufacturer’s specifications. IL-9 levels were outside of the assay’s detection limits and are not shown (Control n=6, K_3_Citrate n =6-7). Sample size varied between assays based on the volume of plasma required and the volume of plasma collected from each mouse.

Compounds shown in Figure 2 I-AA were measured using NMR at LabCorp (Control n=6, K_3_Citrate n =7). NMR spectra were acquired on a Vantera*^®^* Clinical Analyzer, a 400 MHz NMR instrument, from EDTA plasma samples as described for the *NMR LipoProfile^®^* test (Labcorp, Morrisville, NC)(80,81). The *NMR MetaboProfile* analysis, using the LP4 lipoprotein profile deconvolution algorithm, reports lipoprotein particle concentrations and sizes, as well as concentrations of metabolites such as total branched chain amino acids, valine, leucine, and isoleucine, alanine, glucose, citrate, total ketone bodies, β-hydroxybutyrate, acetoacetate, acetone. The diameters of the various lipoprotein classes and subclasses are: total triglyceride-rich lipoprotein particles (TRL-P) (24-240 nm), very large TRL-P (90-240 nm), large TRL-P (50-89 nm), medium TRL-P (37-49 nm), small TRL-P (30-36 nm), very small TRL-P (24-29 nm), total low density lipoprotein particles (LDL-P) (19-23 nm), large LDL-P (21.5-23 nm), medium LDL-P (20.5-21.4 nm), small LDL-P (19-20.4 nm), total high density lipoprotein particles (HDL-P) (7.4-13.0 nm), large HDL-P (10.3-13.0 nm), medium HDL-P (8.7-9.5 nm), and small HDL-P (7.4-7.8 nm). Mean TRL, LDL and HDL particle sizes are weighted averages derived from the sum of the diameters of each of the subclasses multiplied by the relative mass percentage. Linear regression against serum lipids measured chemically in a apparently healthy study population (n=698) provided the conversion factors to generate NMR-derived concentrations of total cholesterol (TC), triglycerides (TG), TRL-TG, TRL-C, LDL-C and HDL-C. NMR-derived concentrations of these parameters are highly correlated (r ≥0.95) with those measured by standard chemistry methods. Details regarding the performance of the assays that quantify BCAA, alanine and ketone bodies have been reported(82,83). While these NMR assays have been analytically validated for use with human specimens, full analytical validation studies have not been performed in rodent specimens.

### Tissue Processing and Ex Vivo Micro-Computed Tomography

Caudal spine segments Ca6-Ca8 (n=7 mice/treatment; 2 discs, 1 vertebrae/mouse; 14 discs, 7 vertebrae/treatment) were dissected and immediately fixed in 4% PFA in PBS at 4°C for 48 hours. Caudal spine segments Ca8-Ca10 (n=7 mice/treatment; 2 discs, 1 vertebrae/mouse; 14 discs, 7 vertebrae/treatment) were fixed for 2 hours in 4% PFA in PBS at 4°C. Following fixation, μCT scans (Bruker Skyscan 1275; Bruker, Kontich, Belgium) were performed on all motion segments. An aluminum filter was used; all scans were conducted at 50 kV and 200 μA, with an exposure time of 85 ms, yielding a resolution of 8 μm. Three-dimensional image reconstructions were generated in nRecon (Bruker), analyzed in CTan (Bruker), and visualized using CTan and CTVox (Bruker). Size, trabecular, cortical, and mineral density parameters were analyzed according to previously reported methods(78,84).

### FTIR

Ca8-Ca10 motion segments (n=7 mice/treatment) were treated with 30% sucrose, OCT-embedded, and snap-frozen. Cryosections of 10 μm were cut and the Spectrum Spotlight 400 FT-IR Imaging system (Perkin Elmer) was used to collect IR spectral imaging data in the mid-IR region from 4,000–750/cm at 8/cm spectral resolution and 25 μm spatial resolution. Absorbance for the amide I region (1665 cm^-1^), collagen side chain vibrations (1338 cm^-1^), phosphate vibration region (960 cm^-1^), and carbonate (870 cm^-1^) were recorded(85). Spectra were processed, and images were generated using ISys Chemical Imaging Analysis software v. 5.0.0.14 (Malvern Panalytical Ltd). To remove noise, spectra underwent a baseline subtraction, followed by normalization and spectral subtraction of the 1736 cm^-1^ peak, which results from the cryotape used to mount calcified sections. Reported spectra and images reflect these corrections. Plotted data reflect all mineralized discs in the Ca8-Ca10 region from Control (n = 12) and K_3_Citrate (n=5) mice.

### Spinal Tissue Processing and Histology

After μCT was completed, Ca6-Ca8 motion segments (n=7 mice/treatment) underwent 21 days of decalcification in 20% EDTA at 4°C, followed by paraffin embedding. Coronal sections of 7 μm were generated, and histoclear deparaffinization followed by graded ethanol rehydration preceded all staining protocols. Safranin O/Fast Green/Hematoxylin staining was conducted and visualized using 5x/0.15 N-Achroplan and 20x/0,5 EC Plan-Neofluar (Carl Zeiss) objectives on an AxioImager 2 microscope and Zen2™ software (Carl Zeiss Microscopy). This staining was used to evaluate disc structure, and four blinded graders scored NP and AF compartments using Modified Thompson Grading (63,86). Picrosirius red staining was conducted and imaged in the brightfield and under polarized light using 4x Pol/WD 7.0 objectives on an Eclipse LV100 POL microscope (Nikon). NIS Elements Viewer software (Nikon) was then used to evaluate the areas of the disc occupied by green, yellow, or red pixels. For all immunohistochemical stains, antibody-specific antigen retrieval was conducted by way of incubation in either chondroitinase ABC for 30 minutes at 37°C or proteinase K for 8 minutes at room temperature. Sections were then blocked in 5-10% normal serum in PBS-T (0.4% Triton X-100 in PBS) and incubated overnight with primary antibodies detailed in Supplementary File 2.1. Tissue sections were washed with PBS-T and incubated in the dark with the appropriate Alexa Fluor® -594 or -647 conjugated secondary antibody (1:700; Jackson ImmunoResearch Laboratories, Inc.) for one hour at room temperature. All stained sections were washed with PBS-T and mounted with ProLong™ Diamond Antifade Mountant with DAPI (Fisher Scientific, P36971). Stains were visualized with an AxioImager 2 (Carl Zeiss Microscopy), using 5x/0.15 N-Achroplan and 20x/0,5 EC Plan-Neofluar objectives, an X-Cite® 120Q Excitation Light Source (Excelitas Technologies), AxioCam MRm camera (Carl Zeiss Microscopy), and Zen2TM software (Carl Zeiss Microscopy). Exposure settings remained constant across treatments for each stain (n=7 mice/treatment/stain, 2 discs/mouse, 14 discs/treatment/stain).

### Knee Histology and Histomorphometry Analysis

Hindlimbs were fixed and scanned for μCT according to the previously described methods(45). 3D reconstructions were evaluated to count the calcification nodules present in each joint(35). Tissues were then decalcified in 20% EDTA at 4°C for 21 days, followed by paraffin embedding. Tissue sections were cut at 5mm in the coronal plane and stained with hematoxylin and eosin (H&E) or toluidine blue and OA severity was analyzed by Articular Cartilage Structure (ACS), toluidine blue, osteophyte, and synovial hyperplasia scoring(44,45). Histomorphometric analysis of articular cartilage thickness and area, calcified cartilage thickness and area, and subchondral bone thickness and area were analyzed according to previous documentation(45).

### RNA Collection and Isolation

Caudal NP tissues from control and K_3_Citrate cohorts (n=4 mice/cohort) were micro-dissected and immediately placed in RNAlater® Reagent (Invitrogen, Carlsbad, CA). Tissues were stored at -80°C until RNA was extracted from the lysates using the RNeasy® Mini kit (Qiagen).

### RNA-Sequencing and Bioinformatic Analysis

Libraries for whole transcriptome RNA sequencing were prepared using the Stranded Total RNAseq with Ribo-zero Plus kit (Illumina, San Diego, CA) as per manufacturer’s instructions starting with an input of 50 ng of RNA and 14 cycles of final PCR amplification. Library size was assessed using the 4200 TapeStation and the DNA D5000 ScreenTape assay (Agilent, Santa Clara, CA). Library concentration was determined using the Qubit Fluorometer 2.0 (ThermoFisher Scientific, Waltham, MA) as well as by quantitative PCR (KAPA Biosystems, Wilmington, MA, USA). Sequencing was conducted using GENEWIZ® NGS Services from Azenta Life Sciences (South Plainfield, NJ, USA). Libraries were multiplexed and clustered onto a flow cell. After clustering, the flow cell was loaded onto the NovaSeq 6000 or equivalent instrument according to manufacturer’s instructions. The samples were sequenced using a Paired End (PE) 100 x 10 x 10 x 10 x 100 configuration and 1% PhiX spike-in. Raw sequence data (.bcl files) generated from Illumina NovaSeq was converted into FASTQ files and de-multiplexed using Illumina bcl2fastq 2.20 software. One mis-match was allowed for index sequence identification. Sequence reads were aligned to the mm10 genome build using STAR 2.7.11b, and counts were retrieve with quantMode. RNA-seq raw counts and TPM are detailed in Supplementary File 1.1-1.3. Data are deposited in the NCBI GEO database under the accession ID GSE270561.

DEGs were analyzed using the GTAC-CompBio Analysis Tool (PercayAI Inc., St. Louis, MO). CompBio performs a literature analysis to identify relevant biological processes and pathways represented by the input differentially expressed entities, in this case, DEGs(78,87). Conditional probability analysis is utilized to compute the statistical enrichment of biological concepts (processes/pathways) over those that occur by random sampling. Related concepts built from the list of differentially expressed entities are further clustered into higher-level themes (e.g., biological pathways/ processes, cell types, and structures, etc.). Within CompBio, scoring of entity (DEG), concept, and overall theme enrichment is accomplished using a multi-component function referred to as the Normalized Enrichment Score (NES). Compbio outputs resulting from downregulated and upregulated DEG analysis are detailed in Supplementary File 1.4-1.5.

### Digital Image Analysis

All immunohistochemical quantification was conducted in greyscale using the Fiji package of ImageJ(88). Images were thresholded to create binary images, and NP, AF, and subchondral bone regions were manually defined using the Freehand Tool. These defined regions of interest were then analyzed either using the Area Fraction measurement or Analyze Particles (TUNEL and cell number quantification) functions.

### ATDC5 Cell Culture

Chondrogenic ATDC5 mouse cells were cultured and differentiated according to the protocol established by Newton et al(48). Briefly, cells were cultured in differentiation medium comprised of DMEM/F-12 with GlutamAX I (Gibco™, 10565018), 5% FBS, 1% Insulin-Transferrin-Selenium-Sodium Pyruvate (ITS-A) (Gibco™, 51300044), and 2% Penicillin-Streptomycin (Corning™, 30001CI) at a density of 4,000 cells/cm^2^ in multi-well plates. Media was changed every 2-3 days, and after 6 days, when the cells reached confluency, treatment-specific media supplementation began. CT cells continued in the previously described differentiation medium. Diff. CT were supplemented with 10mM β-Glycerophosphate (Sigma-Aldrich, G9422) and 50 μg/ml L-ascorbate-2-phosphate. Diff. + 0.25 and Diff + 0.50 treatment groups received 0.25 mM and 0.50 mM K_3_Citrate (Sigma-Aldrich, C3029), respectively. The cultures were continued until day 7, 10, 14, or 21, depending on the subsequent experiment.

### Alizarin Red Staining and Quantification

Alizarin red staining was conducted according to a standardized protocol. Cells were rinsed with PBS and fixed with 4% PFA for 1 hour at room temperature. Cells were then washed with PFA, incubated with 2% (w/v) Alizarin Red (pH 4.1-4.3) for one hour at room temperature on a gentle rocker, and washed with water. To quantify the stain, 10% acetic acid was added to each well of the culture plate and incubated for 30 minutes, with shaking. Resulting solutions were scraped from the culture plates, transferred to microfuge tubes, vortexed, and heated at 85°C for 10 minutes. Hot tubes were then placed in ice for 5 minutes, and the slurry was centrifuged at 20,000 rcf for 15 minutes. The supernatant was then brought to pH 4.1-4.5 with 10mM sodium hydroxide, and optical density of the resulting solution was read for each sample.

### ATDC5 RNA Isolation and qRT-PCR

RNA was extracted from ATDC5 cells according to manufacturer’s protocol, using an RNeasy® Mini kit (Qiagen, 74104), and this RNA was converted to cDNA using EcoDryTM Premix (Clontech Laboratories, 639548). Template cDNA and gene-specific primers were combined with SYBR Green master mix (Applied Biosystems, A25742) and mRNA expression was quantified using the QuantStudio™ 3 System (Applied Biosystems). Gene expression was normalized to Hprt. Primers were synthesized by Integrated DNA Technologies and are listed in Supplementary File 2.2.

### Seahorse Metabolic Analyses

Three assays were conducted using a Seahorse SF Analyzer (Agilent): glycolytic capacity, ATP production, and MitoStress. For all assays, ATDC5 cells were plated in a 24-well Seahorse XF24 V7 PS microplate (Agilent, 100777-004) and cultured according the methods described under *ATDC5 Cell Culture* for Diff CT, Diff. + 0.25, and Diff. + 0.50 conditions until either 7 or 14 days. On the day of the assay, media was removed from the cells, and they were washed 3 times with 500 μL of Krebs Ringer Phosphate HEPES (KRPH) and incubated at 37°C for 1 hour without CO_2_; for the MitoStress test, cells were incubated in the KPRH buffer plus their relevant substrates (5 mM glucose, 5 mM glucose + 0.25 mM K3Citrate, or 5mM glucose + 0.50 mM K_3_Citrate :: Diff. CT, Diff. + 0.25, and Diff. + 0.50). The output for all Seahorse assays were OCR and ECAR. To evaluate glycolytic capacity, the methodology detailed by Mookerjee et al. was used(50). Injections throughout the assay were as follows: 1) Substrate (5 mM glucose, 5 mM glucose + 0.25 mM K3Citrate, or 5mM glucose + 0.50 mM K_3_Citrate :: Diff. CT, Diff. + 0.25, and Diff. + 0.50), 2) 1 μM Rotenone and 1 μM Myxothiozol (for all treatment groups), and 3) 200 μM Monensin and 1 μM FCCP (for all treatment groups). Glycolytic and oxidative ATP production were measured and calculated according to the methodology developed by Mookerjee et al(51). Injections throughout the assay were as follows: 1) Substrate (5 mM glucose, 5 mM glucose + 0.25 mM K_3_Citrate, or 5mM glucose + 0.50 mM K_3_Citrate :: Diff. CT, Diff. + 0.25, and Diff. + 0.50), 2) 2 μg Oligomycin and 3) 1 μM Rotenone and 1 μM Myxothiozol. The MitoStress test was conducted according to manufacturer’s specifications(54). Injections throughout the assay were as follows: 1) 2 μg Oligomycin, 2) 1 μM FCCP, 3) 1 μM Rotenone and 1 μM Myxothiozol. The rates of oxygen consumption and extracellular acidification were normalized to the protein content of the appropriate well for all assays.

### Statistical Analyses

Statistical analysis was performed using Prism 10 (GraphPad, La Jolla, CA, USA) with data presented as mean ± standard deviation (SD), p<0.05. For in vivo analyses, data distribution was checked with the Shapiro-Wilk normality test; a Student ’st test was applied to normally distributed data, and a Mann Whitney test was applied to non-normally distributed data. Distribution data were compared using a χ^2^ test.

## Funding and Acknowledgments

This study is supported by the grants from NIAMS R01AR055655, R01AR064733, and R01AR074813 to MVR and R01AR082460 to KvdW and MVR.

## Author Contributions

OKO, JAC, KvdW, and MVR designed the project. OKO, JJM, BNK, AS, JAC, QW, MC, and FN performed all experiments and analyzed data. OKO, JAC, KvdW, and MVR wrote and edited the manuscript.

## Data Availability Statement

The RNA-sequencing dataset generated in this study is publicly available in the NCBI GEO database under the accession ID GSE270561.

---

**Supplementary Figure 1.**
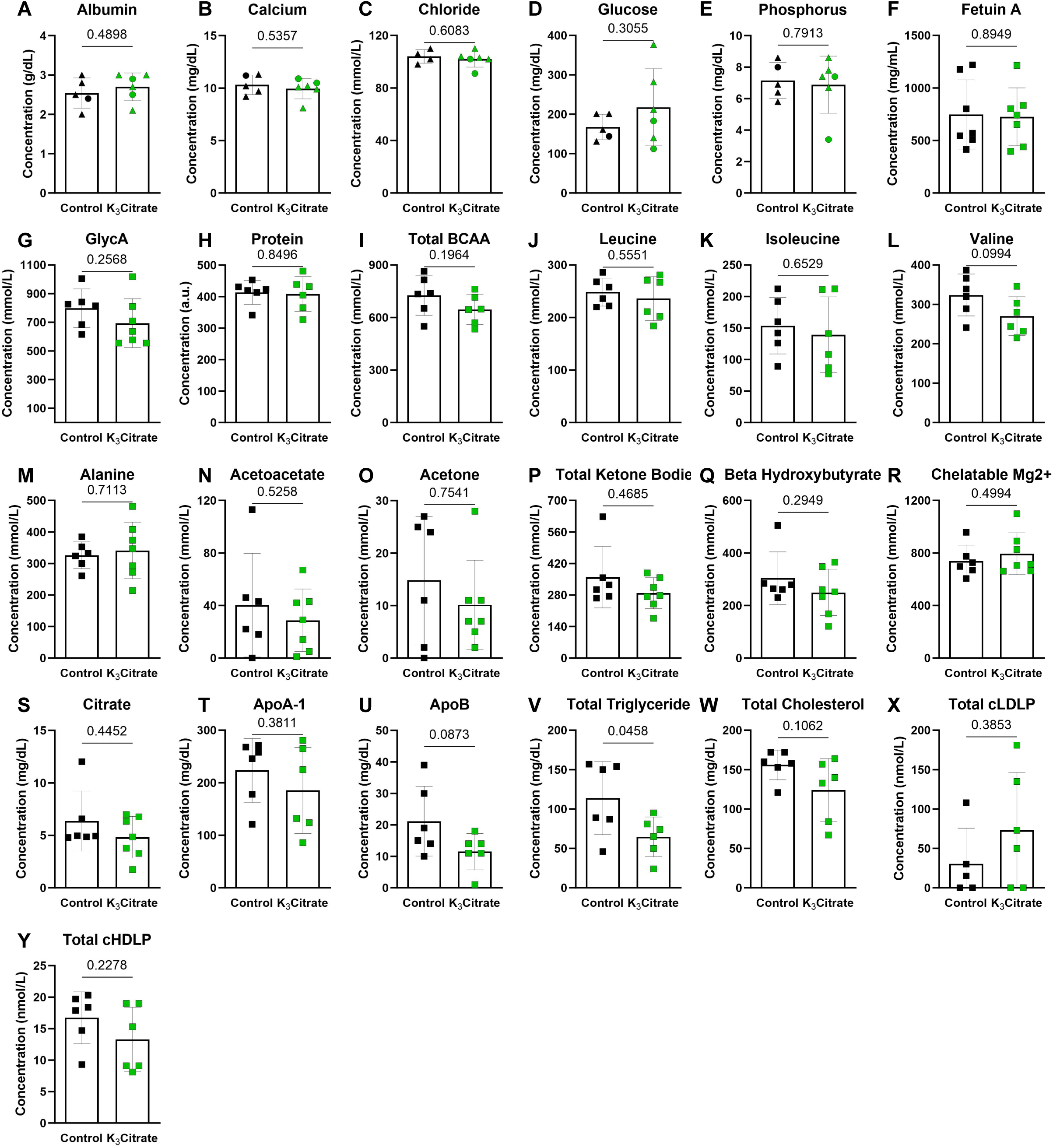
Plasma chemistry is stable with K_3_Citrate supplementation in LG/J mice. **(A)** Albumin, **(B)** Calcium, **(C)** Chloride, **(D)** Glucose, and **(E)** Phosphorus were analyzed using an IDEXX BioAnalytics custom blood chemistry panel (Control: n=5 mice (1F, 4M); K_3_Citrate: n=6 mice (2F, 4M)). **(F)** Fetuin-A, an inhibitor of mineralization, showed no differences across groups (Control: n=7 mice (2F, 5M); K_3_Citrate: n=7 mice (3F, 4M)). **(G)** GlycA, **(H)** Protein, **(I)** total branched chained amino acids (BCAA), **(J)** Leucine, **(K)** Isoleucine, **(L)** Valine, **(M)** Alanine, **(N)** Acetoacetate, **(O)** Acetone, **(P)** Total Ketone Bodies, **(Q)** Beta Hydroxybutyrate, **(R)** Chelatable Mg2+, **(S)** Citrate, **(T)** ApoA-1, **(U)** ApoB, **(V)** Total Trigliyceride, **(W)** Total Cholesterol, **(X)** total calibrated low-density lipoprotein particle (cLDLP), and **(Y)** total calibrated high-density lipoprotein particle (cHDLP) were measured using NMR at LabCorp Global Research Services, and showed no changes with K_3_Citrate supplementation (Control: n=6 mice (1F, 5M); K_3_Citrate: n=7 mice (3F, 4M)). Sample size varied between assays based on the volume of plasma required and the volume of plasma collected from each mouse. Significance was determined using an unpaired t-test or Mann-Whitney test, as appropriate.

**Supplementary Figure 2.**
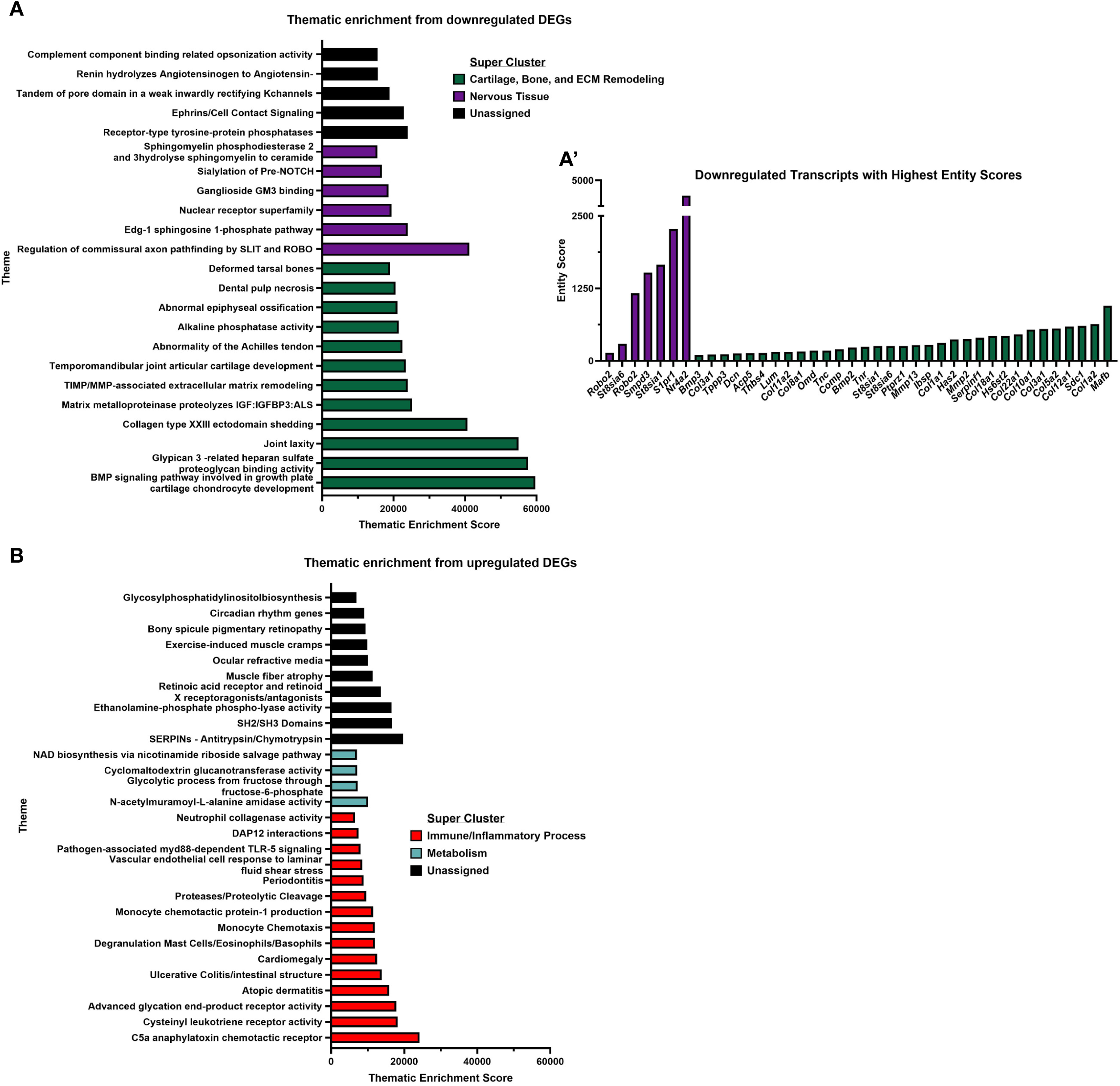
Analysis of upregulated DEGs and most enriched downregulated transcripts. **(A)** RNA-sequencing analysis shows downregulated DEGs have high thematic enrichment. **(A’)** Transcripts within each super cluster that have the highest entity scores (cutoff of 100). **(B)** RNA-sequencing analysis shows upregulated DEGs have high thematic enrichment. (Control: n=4 mice (1F, 3M); K_3_Citrate: n=7 mice (2F, 2M))

**Supplementary Figure 3.**
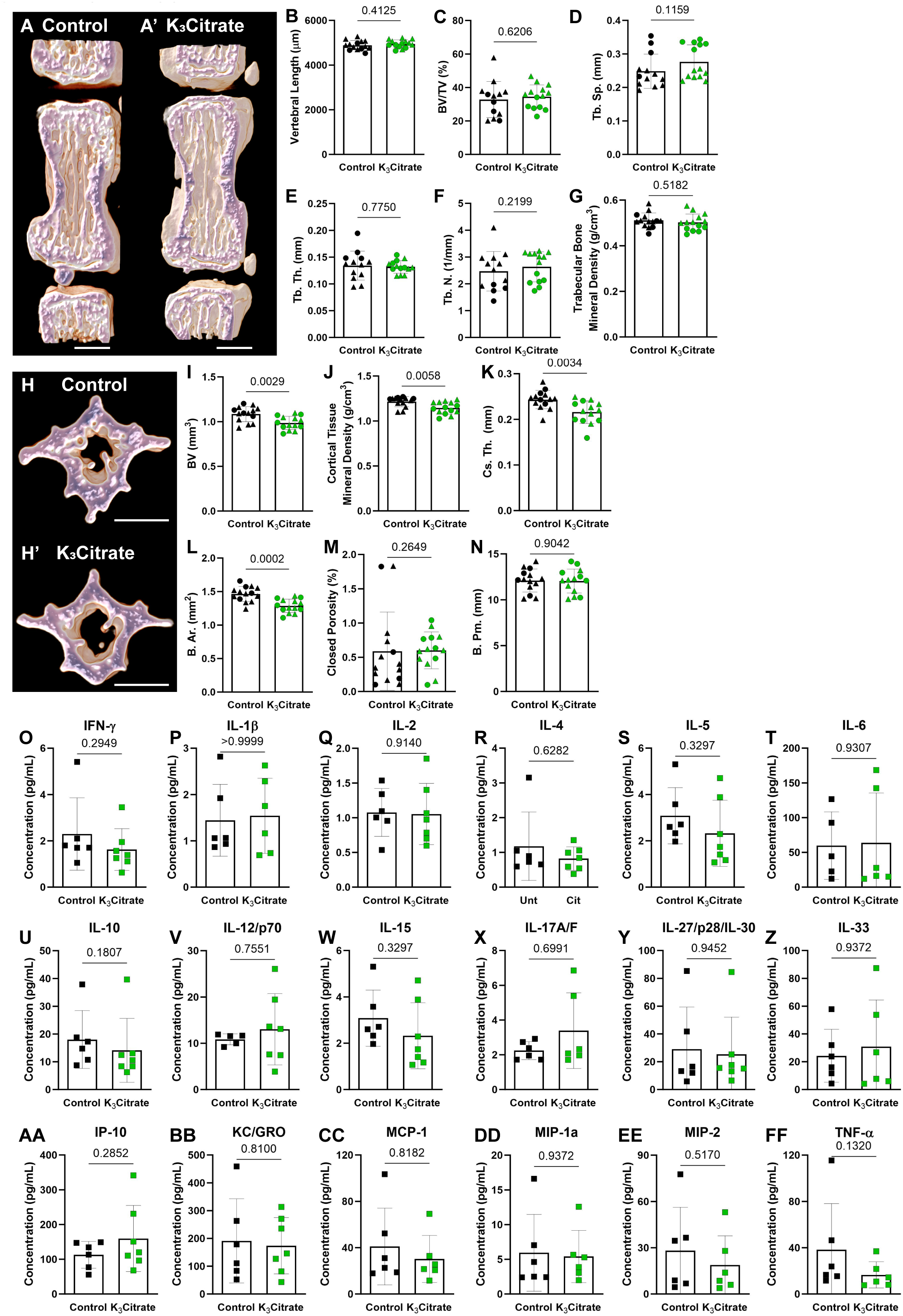
K_3_Citrate supplementation minimally impacts the caudal vertebrae in LG/J mice. (A-A’) Representative μCT reconstructions of the hemi-section caudal motion segments., and **(B)** vertebral length was unchanged. Trabecular properties of **(C)** bone volume fraction (BV/TV), **(D)** trabecular separation (Tb. Sp.), **(E)** trabecular number (Tb. N.), **(F)** trabecular thickness (Tb. Th.), and **(G)** trabecular bone mineral density were unchanged in K_3_Citrate mice. **(H-H’)** Representative μCT reconstructions central cross sections of the caudal vertebrae. Analysis of the cortical properties **(I)** bone volume (BV), **(J)** cortical tissue mineral density, **(K)** cross sectional thickness (Cs. Th.), **(L)** mean cross-sectional bone area (B. Ar.), **(M)** closed porosity, and **(N)** bone perimeter (B. Pm.) revealed cortical thinning of K_3_Citrate caudal vertebrae. (Control mice: n=7 mice (2F, 5M); K_3_Citrate mice: n=7 mice (3F, 4M); 2 vertebrae/mouse; 28 discs, 14 vertebrae/treatment) Data are shown as mean ± SD. Significance was determined using an unpaired t-test or Mann-Whitney test, as appropriate. Multiplex assay analysis showed no significant changes in the plasma concentrations of **(O)** IFN-γ, **(P)** IL-1β, **(Q)** IL-2, **(R)** IL-4, **(S)** IL-5, **(T)** IL-6, **(U)** IL-10, **(V)** IL-12/p70, **(W)** IL-15, **(X)** IL-17A/F, **(Y)** IL-27/p28/IL-30, **(Z)** IL-33, **(AA)** IP-10, **(BB)** KC/GRO, **(CC)** MCP-1, **(DD)** MIP-1α, **(EE)** MIP-2, **(FF)** TNF-α (Control: n=6 mice (1F, 5M); K_3_Citrate: n=7 mice (3F, 4M)). Data are shown as mean ± SD. Significance was determined using an unpaired t-test or Mann-Whitney test, as appropriate.

**Supplementary Figure 4.**
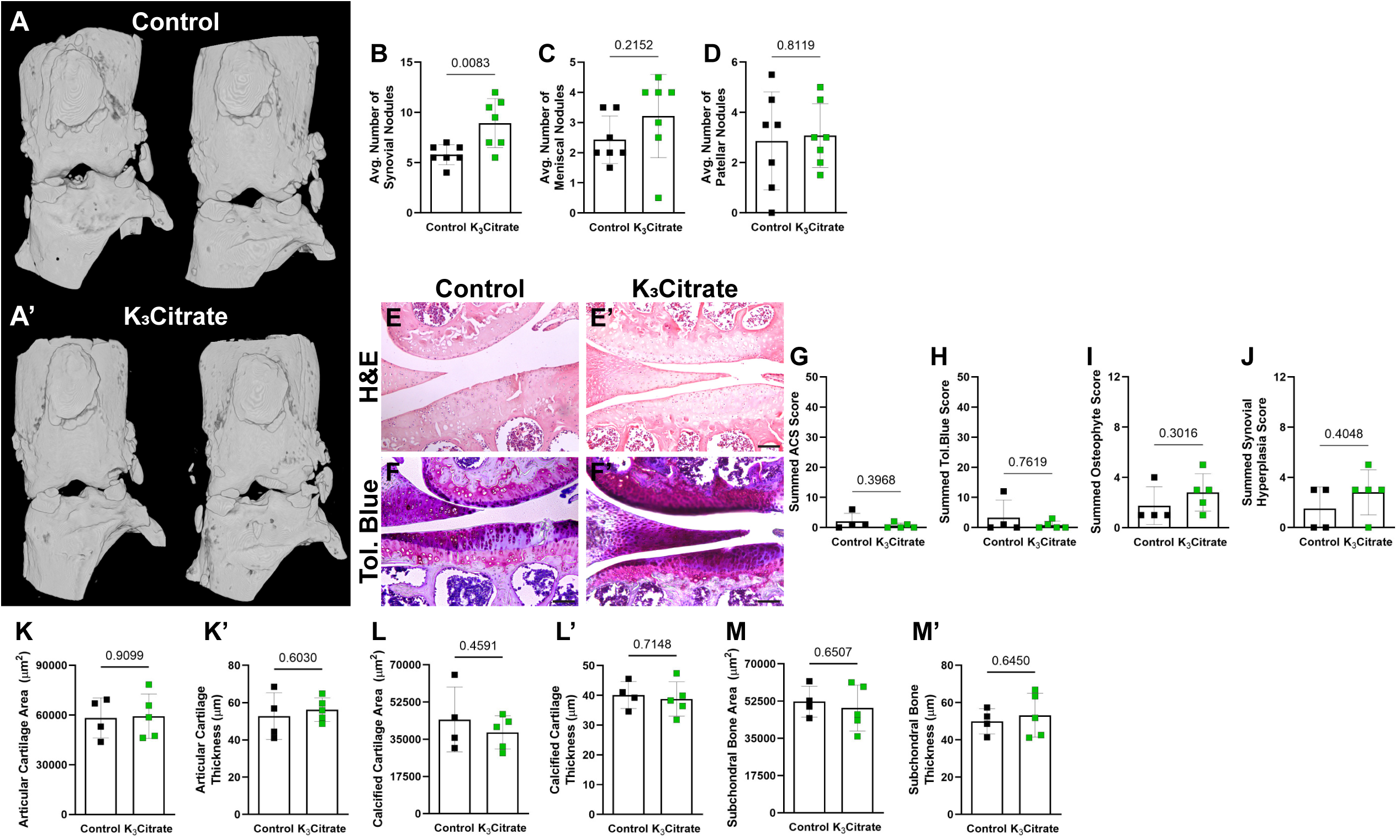
K_3_Citrate alters knee calcification without impact on cartilage and bone morphology. **(A-A’)** Representative μCT reconstructions of the knees of control and K_3_Citrate mice. Quantification of calcification nodules in the **(B)** synovium, **(C)** meniscus, and **(D)** patellar. **(E-E’)** Representative hematoxylin and eosin (H&E) staining and (**F-F’)** toluidine (Tol.) blue staining of the lateral tibial plateau. Summed **(G)** ACS score, **(H)** toluidine blue score, **(I)** osteophyte score, and **(J)** synovial hyperplasia score show no difference between cohorts. Additionally, K_3_Citrate supplementation did not result in changes to the area or thickness of **(K-K’)** articular cartilage, **(L-L’)** calcified cartilage, or **(M-M’)** subchondral bone. (Control: n=4 mice (1F, 3M); K_3_Citrate: n=5 mice (3F, 2M)). Data are shown as mean ± SD. Significance was determined using an unpaired t-test or Mann-Whitney test, as appropriate.

**Supplementary Figure 5.**
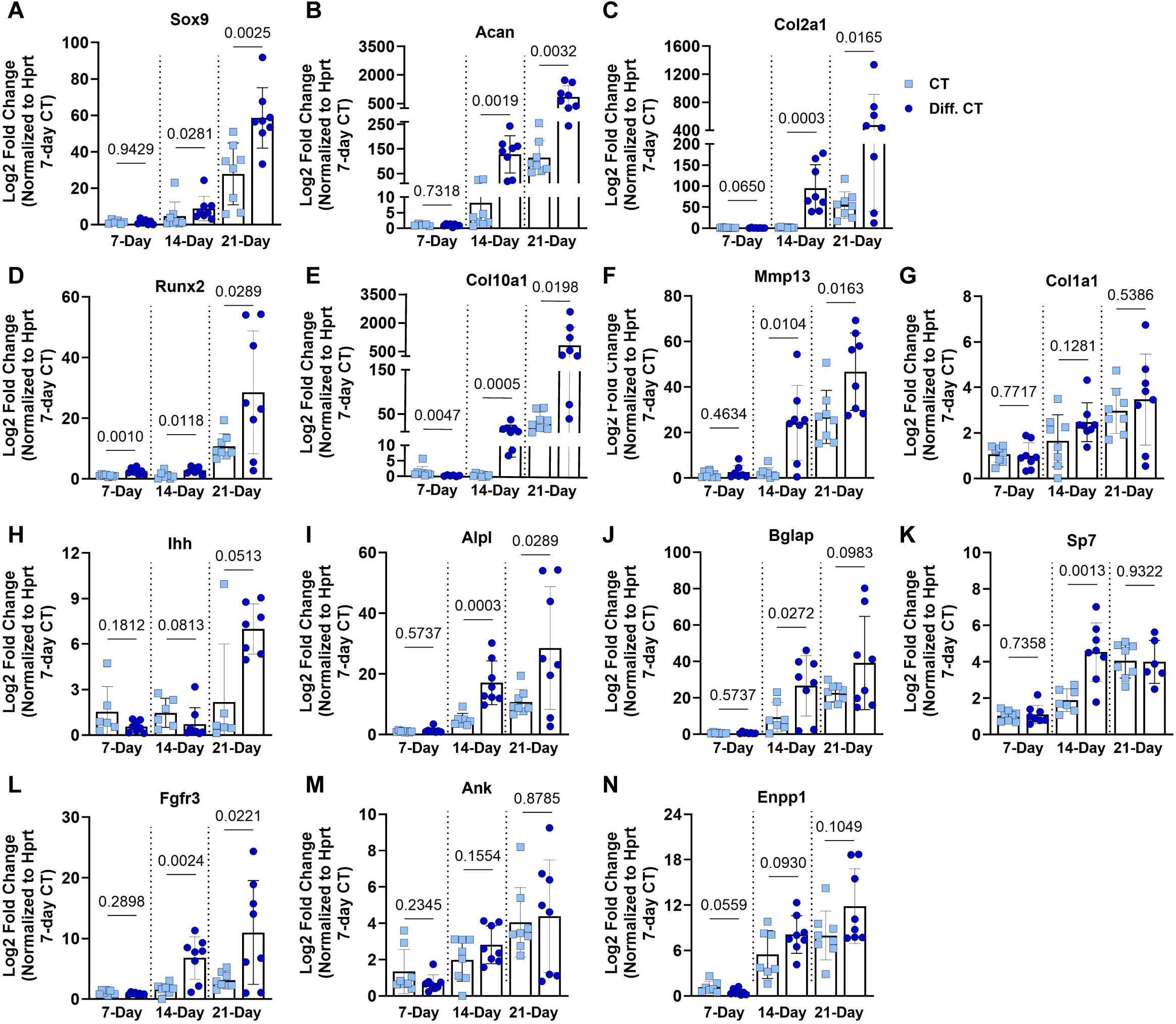
Validation of accelerated differentiation protocol in ATDC5 cells. CT and Diff. CT treatment groups were compared for all chondrogenic markers assessed to verify accelerated differentiation in ATDC5 cells receiving β-glycerophosphate and ascorbic acid: **(A)** Sox9, **(B)** Acan, **(C)** Col2a1, **(D)** Runx2, **(E)** Col10a1, **(F)** Mmp13, **(G)** Col1a1, **(H)** Ihh, **(I)** Alpl, **(J)** Bglap, **(K)** Sp7, and **(L)** Fgfr3, **(M)** Ank, and **(N)** Enpp1. (n=8 sets/timepoint, 2 averaged replicates/set) Data are shown as mean ± SD. Significance was determined using an unpaired t-test or Mann-Whitney test, as appropriate.

**Supplementary Figure 6.**
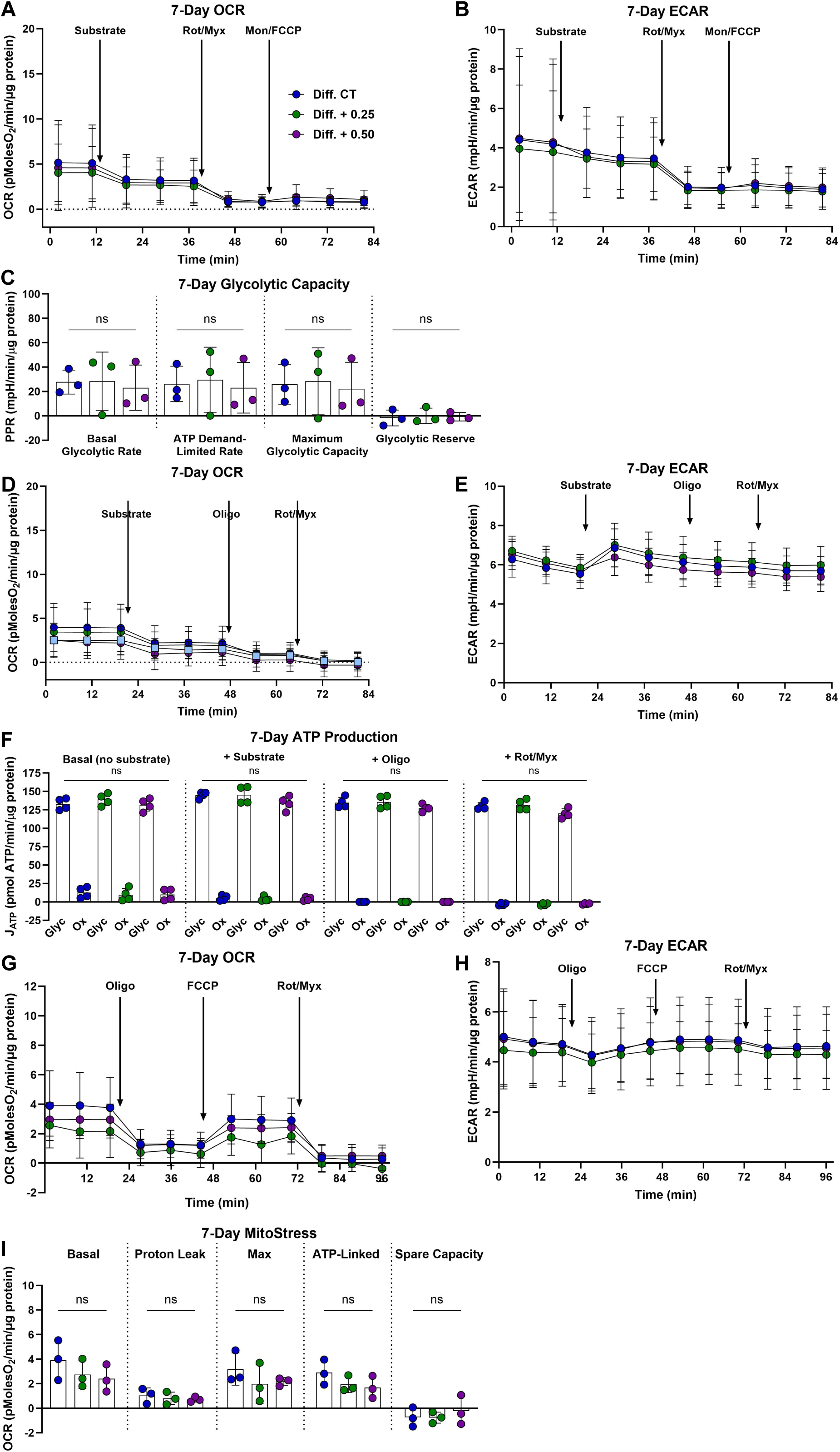
K_3_Citrate supplementation does not impact glycolytic or oxidative metabolism in ATDC5 cells cultured for 7 days. **(A)** OCR and **(B)** ECAR traces for ATDC5 cells cultured with or without K_3_Citrate **(C)** to evaluate glycolytic capacity and glycolytic reserve. **(D)** OCR and **(E)** ECAR traces for ATDC5 cells cultured with or without K_3_Citrate **(F)** to evaluate glycolytic and oxidative ATP production rates. **(G)** OCR and **(H)** ECAR traces for ATDC5 cells cultured with or without K_3_Citrate **(I)** for the classical Mito Stress test to evaluate key parameters of mitochondrial function. (n = 3 sets, 3-4 replicates/set) Data are shown as mean ± SD. Significance was determined using an ANOVA or Kruskal-Wallis test, as appropriate.

