## Supplemental File 2 for "Oral citrate supplementation mitigates age-associated pathological intervertebral disc calcification in LG/J mice"

**Supplementary Tables**

| **Antibody** | **Concentration** | **Company** | **Catalog** |
| --- | --- | --- | --- |
| anti-carbonic anhydrase 3 | 1:150 | Santa Cruz Biotechnology | sc-50715 |
| anti-glucose transporter I | 1:200 | Abcam | ab40084 |
| Vector® Blue Substrate Kit, Alkaline Phosphatase (TNAP) | Manufacturer's Specifications | Vector Laboratories | SK-5300 |
| anti-aggrecan | 1:50 | Millipore Sigma | AB1031 |
| anti-collagen X | 1:500 | Abcam | ab58632 |
| In Situ Cell Death Detection Kit (TUNEL) | Manufacturer's Specifications | Roche Diagnostics | 12156792910 |
| **Supplementary Table 2.1.** Immunohistochemistry reagents | | |  |

| **Gene** | **Forward Sequence (5’-3’)** | **Reverse Sequence (5’-3’)** |
| --- | --- | --- |
| *Hprt1* | CCTCATGGACTGATTATGGACAG | TCAGCAAAGAACTTATAGCCCC |
| *Sox9* | GAAGTCGCTGAAGAACGGACAAG | GCTGTAGTGAGGAAGGTTGAAGGG |
| *Acan* | GATCTACCGCTGTGAAGTGATG | GGGTGTAGCGTGTGGAAATAG |
| *Col2a1* | GCTGGTGAAGAAGGCAAACGAG | CCATCTTGACCTGGGAATCCAC |
| *Runx2* | ACTCTTCTGGAGCCGTTTATG | GTGAATCTGGCCATGTTTGTG |
| *Col10a1* | AACGCCCACAGGCATAAA | GGAATGCCTTGTTCTCCTCTTA |
| *Mmp13* | GACACAGCAAGCCAGAATAAAG | GGAAAGCAGAGAGGGATTAACA |
| *Col1a1* | AGACCTGTGTGTTCCCTACT | GAATCCATCGGTCATGCTCTC |
| *Ihh* | CTCAGACCGTGACCGAAATAAG | TGGGCCTTGGACTCGTAATA |
| *Alpl* | GGAATACGAACTGGATGAGAAGG | GGTTCCAGACATAGTGGGAATG |
| *Bglap* | CTGCCCTAAAGCCAAACTCT | AGCTGCTGTGACATCCATAC |
| *Sp7* | TGGAGAGGGAAAGGGATTCT | GAAATCTACGAGCAAGGTCTCC |
| *Fgfr3* | TCTGGGCTAAGGATGGTACA | ATCTTCGTGGGAGGCATTTAG |
| **Supplementary Table 2.2.** Mouse primer sequences used in ATDC5 experiments. | | |
